# Local Adaptation of Life-History Traits in a Seasonal Environment

**DOI:** 10.1101/2025.02.25.640023

**Authors:** Rebekah Hall, Ailene MacPherson

## Abstract

Populations are often spread across a spatially heterogeneous landscape, connected by migration. Consequently, the question arises whether divergent selective forces created by spatial heterogeneity can overcome the homogenising force of migration and loss of diversity through genetic drift to favour different traits across space. Such spatial heterogeneity in the population due to divergent selection is known as local adaptation. While local adaptation has been studied in a variety of settings, it remains unclear whether local adaptation of certain life-history traits can arise. Life-history traits, such as those determining an organism’s fecundity (the parameter *r*) and ability to compete for resources (the parameter *K*) demonstrate unique eco-evolutionary feedback loops due to their direct relationship to individual fitness. Classic theory holds that in a constant environment, evolution maximises the population’s competitive ability. Divergent selective pressures on life-history traits requires complex environmental differences, such as heterogeneous patterns of seasonality. We consider life-history evolution in a Lotka-Volterra model with three types of seasonal perturbations: repeated, sudden crashes in population size, fluctuating death rates, and fluctuating resource levels. We show that fluctuating resources cannot change the evolutionary outcome, but that sufficiently harsh population crashes or fluctuating death rates favour increased fecundity over competitive ability. Finally, we apply deterministic and stochastic modelling to study local adaptation of an island population to periodic population crashes in an island-mainland model. We find that local adaptation favouring *r*-selected individuals again arises when conditions are sufficiently harsh, but not so harsh that the island population cannot be sustained in the absence of migration.

## Introduction

Local adaptation describes when populations are heterogeneous across space due to environmental heterogeneity favouring different traits. As a result of divergent selection opposing the homogenising effects of migration and loss of diversity through drift (Dittmar and Schemske, 2023; Haldane, 1930; Wright, 1931; Yeaman and Otto, 2011), local adaptation can help maintain biodiversity and generate novel diversity as the first step in ecological speciation (Schluter, 2001). A population’s ability to adapt to new environmental conditions, as indicated by local adaptation, is an important determinant in its ability to colonise and persist in novel habitats and hence during range expansion (Andrade-Restrepo et al., 2019; Gilbert et al., 2017). As organisms adapt to their local environment, they generally experience a fitness cost in other environments (Kawecki and Ebert, 2004; Whitlock, 2015). Hence, the extent of local adaptation is often measured through the comparison of an individual’s fitness in its local environment with the fitness of foreign individuals in that environment (Blanquart et al., 2013; Kawecki and Ebert, 2004).

While most of our understanding of local adaptation is in temporally-constant environments, environments are not only heterogeneous through space, but through time, frequently characterised by both random noise and regular, periodic variation. It is more important now than ever to understand the impacts of temporal environmental variation on evolution and adaptation to local environmental conditions, particuarly in light of climate change. Not only are extremes of seasonal variations increasing with the changing climate (Easterling et al., 2000; Schär et al., 2004), but species are being pressured to disperse into new environments with different environmental conditions (Hastings et al., 2005; Hill et al., 1999; Parmesan and Yohe, 2003). One method for studying patterns of local adaptation in a population undergoing range expansion is through an island-mainland model, in which an established mainland population immigrates to an island with different environmental conditions. The study of local adaptation to temporal variations leads naturally to the study of local adaptation of life-history traits, as such temporal variations are known to influence the evolution of these traits through ecoevolutionary feedbacks (Kremer and Klausmeier, 2013; Lande et al., 2009, 2017; Roughgarden, 1971). To understand how life-history traits can evolve in response to local temporal environmental variation in spatially structured environments we first must consider the evolution of these traits in specific environmental contexts in the absence of gene-flow.

Life-history theory aims to understand the link between ecology and evolution by examining how traits which determine demography—that is, those relating to reproduction, growth, and mortality—evolve under specific environmental conditions and how this gives rise to the observed diversity of life-history traits across environments, species (Stearns, 2000), and here, space. Life-history traits demonstrate unique ecoevolutionary feedback loops due to their direct relationship to individual fitness. They are frequently subject to trade-offs (Mojica et al., 2012; Mérot et al., 2018, 2020; Troth et al., 2018), which are often conceived of within the framework of *r*- and *K*-selection, where *r* and *K* refer respectively to the intrinsic growth rate and carrying capacity in the logistic growth equation (MacArthur and Wilson, 1967; Mueller et al., 1991; Pianka, 1970). These two life-history strategies show contrasting patterns of density dependent selection; individuals with high reproductive rates (a high-*r* life-history strategy) experience a relative fitness advantage at low-population densities that decreases with increas-ing density (negative density dependence), whereas individuals with high competitive abilities (a high-*K* strategy) experience a relative fitness advantage at high population densities that increases with increasing density (positive density-dependence) (Mueller et al., 1991; Roughgarden, 1971). As a result of this density-dependence, theoretical results (Kremer and Klausmeier, 2013; Lande et al., 2009; Roughgarden, 1971) and empirical observations (Mérot et al., 2020; Teuschl et al., 2007; Vasi et al., 1994) suggest that large disruptions to population size are necessary for *r*-selection to occur.

Fluctuations in population sizes, however, will not only mediate the selection on life history traits but the strength of genetic drift, particularly in small range-edge populations or populations at risk of extirpation. In these small populations, drift may dominate over selective forces, leading to the maladapted trait dominating the population. This effect may be exacerbated by a constant influx of maladapted individuals through migration. In the study of local adaptation of life-history traits to seasonal environments, it is therefore unclear whether a regime exists under which conditions are sufficient for *r*-selection to arise, but not so harsh as to allow migration and genetic drift to take over. A comprehensive understanding of local adaptation of life-history traits hence requires the examination of migration-selection-drift balance. The relative importance of genetic drift versus selection is captured by the product of the selection coefficient and the harmonic mean population size through time (Hartl and Clark, 2007; Kimura, 1983; Ohta and Tachida, 1990), the latter of which is disproportionately affected by bottlenecks such as those that may be caused by large seasonal disruptions.

Classic theory holds that in a constant environment, long-term evolution will maximise competitive ability even at the cost of fecundity (MacArthur, 1962). However, theoretical work has shown cases where temporal variations can change this outcome.

Using the *r*-vs. *K*-selection dichotomy, Lande et al. (2009) showed that sufficient environmental noise would lead to evolution maximising fecundity. This result was later extended to an age-structured population (Lande et al., 2017). Roughgarden (1971) used a discrete-time model to consider regular population crashes in the context of overwintering, where there is a fixed number of individuals who survive the winter season. They showed that such seasonal forcings were sufficient for *r*-selected strategies to persist in a population, either coexisting with *K*-selected strategies or taking over the population depending on the degree of (over)dominance in the diploid population. Kremer and Klausmeier (2013) modelled a population with a continuous-time life history in an environment which experiences a “growing” season and a “not-growing” season, as represented by a periodically-changing function for resource levels. They showed that coexistence could arise between *r* and *K* strategies, depending on the the nature of the trade-offs between life-history strategies, the speed of life-history evolution, the length of the period, and the relative lengths of the good versus bad season. Their results also suggested that other ecological mechanisms such as immigration may be important in determining coexistence.

These models cover only a small number of ways in which temporal disruptions may change the evolution of organisms. Other disruptions may include direct fluctuations in population density, or indirect fluctuations mediated by changing resource levels and density-dependent mortality. The nature of these disruptions also interacts with the nature of an organism’s life cycle and whether that cycle is discrete or continuous. How the evolution and local adaptation of life-history traits are affected by this range of different disruptions is still being elucidated. Making predictions for how diverse life-history strategies can be maintained across space requires us to understand both what ‘types’ (e.g., changes in resources, vital rates, or population size) and ‘shapes’ (e.g., severity through time) of environmental disruptions favour high-*r* strategies.

In this paper, we seek to understand how local seasonal disruptions can give rise to local adaptation of life-history traits and hence the spatial coexistence of life-history strategies. Given that *K*-selected strategies are naturally favoured in constant environments, local adaptation arises only when local environmental disruptions are sufficient to favour *r*-selected strategies in the face of ongoing gene flow and drift. To formulate predictions for when local adaptation should arise, we first consider a single population and analyse under what conditions an *r*-selected strategy can evolve as a result of three types of seasonal disruptions: recurring population crashes, a continuously-fluctuating death rate, or a continuously-fluctuating resource supply. Given these single-population results, we then consider local adaptation to a seasonally-varying island environment in the face of ongoing gene flow from a large mainland population with a constant environment. We compare analytical predictions assuming a large but variable population to results from individual-based simulations for small populations experiencing significant genetic drift. Finally, we examine the implications of these results for a wide range of biological systems and suggest future directions for modelling adaptation to temporally-heterogeneous environments.

### Single-population models

To understand under what conditions local adaptation of life-history traits might arise, we first seek to characterise which environmental conditions will favour higher fecundity. Therefore, we begin by considering the evolution of *r* vs. *K* life-history traits in a single population. Evolution is described by a continuous-time Lotka-Volterra model.

While this model is traditionally used as an ecological model of competition between species, such a competition model can equivalently be applied to describe competition between individuals carrying a high-*r* allele (with population density *N*_*r*_) and individuals carrying a high-*K* allele (with population density *N*_*K*_) in a haploid population with asexual reproduction and random mating (Yi and Dean, 2013). Hence, the population dynamics are given by the system of differential equations:

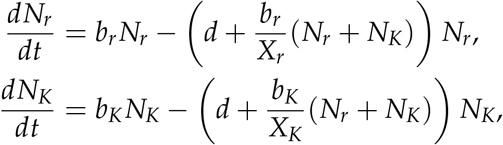

where *b*_*i*_ is the per-capita birth rate, *d* is the death rate, and *X*_*i*_ is the ‘growth limit’ of individuals carrying the *i* = *{r, K}* allele. The growth limit describes the individual’s competitive ability relative to the resources available in the environment by mediating the severity of density-dependent mortality and is linearly related to the individual’s carrying capacity *K*_*i*_ (see Supplementary Material S1). This means that our model assumes a linear increase in deaths as population density and competition increases. We choose parameters to enforce a trade-off such that the high-*r* allele confers a lower carrying capacity 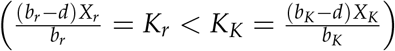,but a higher intrinsic growth rate (*b*_*r*_ > *b*_*K*_). The death rate (*d*) remains constant across both alleles and is assumed to be smaller than both birth rates (see Supplementary Material S1). Note that for any fixed choice of *b*_*i*_ and *X*_*i*_, if the death rate is above a certain threshold, the allele with the higher birth rate will also have the greater carrying capacity. Thus, although there remains a trade-off between the birth rate and the growth limit, there is no longer a trade-off between *r* and *K*. Hence the high-*r* will have the greater fitness in all environmental conditions. In our analytical analysis, we assume a value of *d* that is below this threshold, but we examine a larger range of death rates graphically (Fig. 1C).

**Figure 1:**
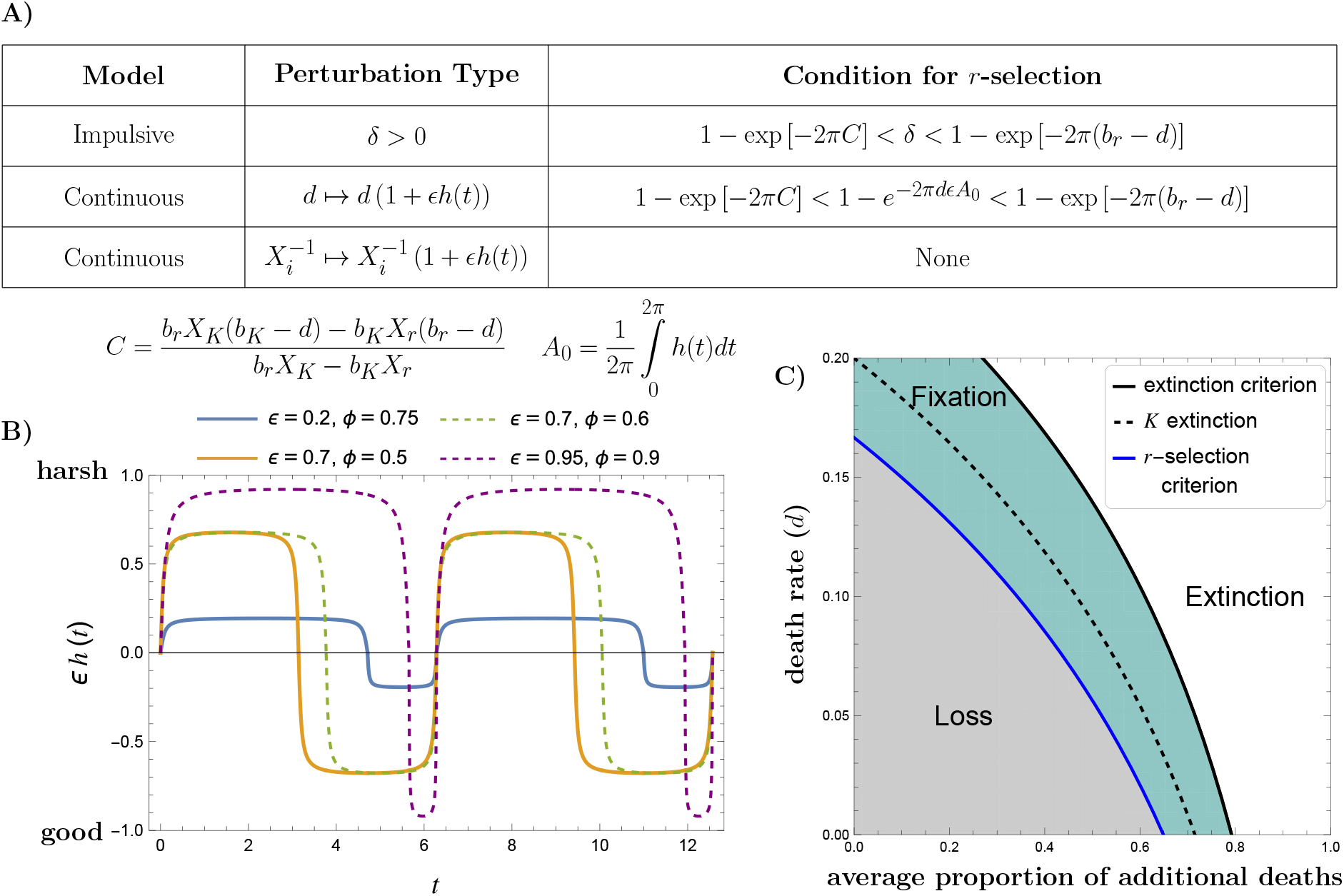
Summary of single population results. (A) We consider three types of seasonal perturbations: a proportion of individuals removed from the population at the end of a season (*δ*); fluctuations in the death rate (*d*); and fluctuations in the growth limit, or available resources (*X*_*i*_). The death rate and growth limit are varied seasonally by incorporating a periodic function, *ϵh*(*t*), which creates seasons with good conditions and seasons with harsh conditions. (B) Example functions for environmental perturbations. The amplitude of the perturbations are determined by *ϵ*, while *ϕ* is the proportion of time spent in the ‘bad’ season. (C) Plot of *r*-selection conditions for perturbation to the average proportion of additional deaths in one season, given by *δ* in the impulsive case and 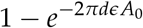 in the continuous case. The high-*r* allele may be lost (grey), may fix (turquoise), or the population may go extinct (white). The *K* extinction line marks the maximum amount of additional death such that a population of exclusively high-*K* allele individuals is viable. Parameters were as follows: *b*_*r*_ = 0.25, *b*_*K*_ = 0.2, *X*_*r*_ = 5000, *X*_*K*_ = 10000.

To maintain consistency with our subsequent model of local adaptation, it is convenient to re-parametrise these two equations in terms of the frequency of the high-*r* allele, *p*, and the total population size, *N*. Hence changes in the allele frequency and changes in the total population size can be described by the following differential equations:

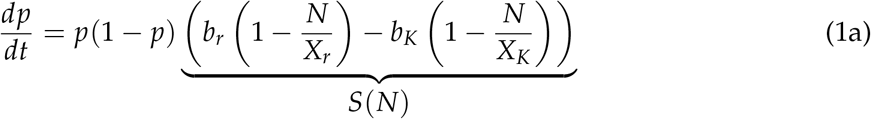

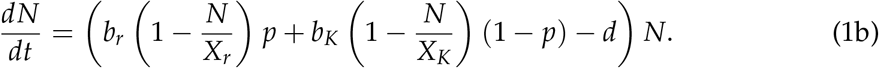

This re-parameterisation additionally allows us to identify the selection coefficient, *S*(*N*), which governs selection on the high-*r* allele. When the selection coefficient is positive, selection favours the high-*r* allele, while it is selected against when the selection coefficient is negative. *S*(*N*) will decrease linearly with increasing population size.

System 1 exhibits three equilibria, describing three possible evolutionary outcomes. The evolution of the population is determined by the stability of these equilibria. Extinction, 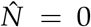,is unstable when *b*_*i*_ > *d* (a condition which is assumed to be met throughout). Fixation of the high-*r* allele, 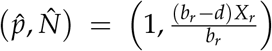,is always unstable, while its loss 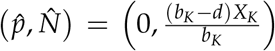,is always stable. Hence the population is always viable, but the high-*r* allele is always lost. This agrees with classic theory which holds that in a constant environment, the high-*K* allele will out-compete the high-*r* allele.

#### Case 1: Seasonal population crashes

In order to incorporate seasonal disruptions, we first consider a case where the population undergoes sudden, periodic population crashes. For example, spikes in mortality due to tidal disruptions and frost events have been observed to favour *K*-selected phenotypes (Mérot et al., 2020; Teuschl et al., 2007). In such cases, seasonal variability comes in the form of population crashes as in the discrete-time model by Roughgarden (1971), but reproduction may not necessarily occur in the same discrete periods as proposed in this model. Therefore, we seek to understand this form of seasonal disruption within a continuous-time framework.

We model spikes in mortality by introducing the instantaneous removal of a proportion *δ* of the population at the end of each season. Since time can be scaled by the rates of births and deaths, the length of the season is arbitrary. For convenience, we choose a season length of 2*π* to maintain consistency with the continuous fluctuation cases below.

This model can be described by modifying System 1 so that *N*(*t*^+^) = (1 − *δ*)*N*(*t*^−^) and *p*(*t*^+^) = *p*(*t*^−^) at the end of each season, where *N*(*t*^−^) and *p*(*t*^−^) are the population size and allele frequency immediately before the crash and *N*(*t*^+^) and *p*(*t*^+^) are the population size and allele frequency immediately after. Note that *p*(*t*^−^) = *p*(*t*^+^) so that the population crash does not change the allele frequency in the population. Hence, there is no selective element to the disruptions.

As in the case of a constant environment, there are three possible outcomes: extinction, fixation, or loss. To determine which of these outcomes will occur under what circumstances, we must look at the stability of each case. Liu et al. (2007) previously applied Floquet analysis to study the stability of a general Lotka-Volterra system with impulses.

Floquet analysis provides an analogous toolkit to that used for stability analysis of an equilibrium point, now for the stability of equilibrium periodic solutions (Klausmeier, 2008). Extending their results to our model (see Supplementary Material S1.1), we can derive the specific result that the high-*r* allele will fix when:

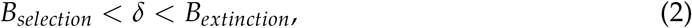

Where:

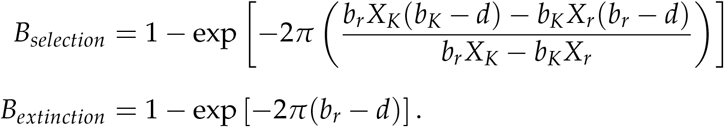

This inequality demonstrates that *r*-selection occurs when population crashes, *δ*, are intermediate between some lower and upper bounds. If *δ* is too small (*δ* < *B*_*selection*_), the disruptions are too mild for selection to favour the high-*r* allele, while if *δ* is too large (*δ* > *B*_*extinction*_), the environment is too harsh to sustain the population and extinction occurs (Fig. 1C). Therefore, we expect that local adaptation may arise in an environment with seasonal population crashes when the extent of the seasonal disruption falls within these bounds. However, it remains to be seen whether the extent of the disruption necessary to favour the high-*r* allele (*B*_*selection*_) is so large that the population is susceptible to swamping by a constant influx of high-*K* alleles, particularly when the effects of gene swamping are combined with genetic drift.

#### Case 2: Continuous seasonality in death rates

Environments may also experience alternating seasons of increased and decreased mortality. For example, the eastern brook trout (*Salvelinus fontinalis*) experiences greater mortality among the youngest age class when temperatures rise, resulting in higher mortality in the summer (Bassar et al., 2016). Predation patterns may also result in seasonal mortality, due to fluctuations in the predator population size or changes in coverage protecting prey (Ferreira and Faria, 2021; Metz et al., 2012).

To model continuous variation in the death rate, we take a similar approach to Rosenblat (1980) and substitute *d* ↦ *d* (1 + *ϵh*(*t*)), where *h*(*t*) is a periodic function and 0 < *ϵ* < 1 is a small quantity, so that fluctuations are small. We could consider a specific periodic function *h*(*t*), such as *h*(*t*) = sin(*t*), but as we will show below, the result is sensitive to the shape of this function. That is, the amount of time spent in the good season (low mortality) as opposed to the bad season (high mortality) is as important to determining the evolutionary outcome as the severity of the seasonal changes. Hence we take a general approach which considers any periodic function. This means that *d* (1 + *ϵh*(*t*)) can describe any seasonal fluctuation of the death rate around the natural death rate *d*. Some examples of possible functions that can be described by our model are pictured in Figure 1B. When *h*(*t*) is positive, the death rate increases and the season is harsh. When *h*(*t*) is negative, the death rate is reduced, corresponding to a good season. By incorporating a periodic perturbation, we must use Floquet analysis (Klausmeier, 2008) to quantify the conditions for our three evolutionary outcomes: fixation, loss, and extinction (see Supplementary Material S1.2). We first examine the conditions for the population to go extinct, finding that extinction occurs if *b*_*r*_ < *d* (1 + *ϵA*_0_), where 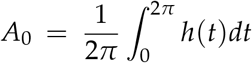 may be interpreted as the average disruption over the course of the season and is determined by both the amplitude and shape of the disruptions. This means that the population goes extinct if the birth rate of the high-*r* allele is less than the average death rate over the course of the season.

Next, we determine the conditions for fixation of the high-*r* allele. We can show that *r*-selection will be favoured when (see Supplementary Material S1.3):

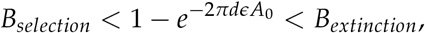

where *B*_*selection*_ and *B*_*extinction*_ are the same bounds as in Equation 2 (Fig. 1A). Like *δ* in Case 1, 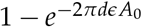 represents the proportion of the population that dies over the course of the season above what would be expected from the natural death rate *d* (Fig. 1C, Supplementary Material S1.3).

Here, the conditions for *r*-selection depend on the strength and shape of the disruption to the death rate. If the amplitude of the disruption is too small (e.g., the blue function in Fig. 1B) or the time spent in the harsh state is too short (e.g., the green function in Fig. 1B), the high-*r* allele will be lost. If fluctuations are too large and the time spent in the harsh state is too long (e.g., the purple function in Fig. 1B), the population will be driven extinct. Hence the high-*r* allele will only fix if the amplitude of the disruption and the time spent in the harsh state are intermediate (e.g., the orange function in Fig. 1B). The effects of the strength and shape of the disruption can be summarised by the average mortality rate over the course of the season. As seen in the analogous inequalities (Fig. 1A), the timing of the additional mortality over the course of the season–whether instantaneous (Case 1) or stretched continuously over the season (Case 2)–does not change when *r*-selection arises. It matters only that there is a sufficiently-high mortality without driving the population extinct. It is therefore in such environments of high mortality that we may expect to see local adaptation, if uninhibited by factors gene swamping and genetic drift.

#### Case 3: Continuous seasonality in resource levels

Seasonality may also disrupt resource availability and hence change the amount of competition and density-dependent mortality. For example, in grasslands, rainfall events influence the quality of vegetation, which in turn determines density-dependent mortality (Branson, 2008). In a chemostat setting, evolution of *Escherichia coli* under alternating periods of feast and famine yielded populations which maximised rapid growth (Vasi et al., 1994).

To model continuous variation in the resource levels, we apply a periodic change to the growth limit by substituting *X*_*i*_ ↦ *X*_*i*_(1 − *ϵh*(*t*)). Here, a positive *h*(*t*) decreases the growth limit. This corresponds to a harsh season with fewer resources. When *h*(*t*) is negative the growth limit increases, representing a good season with more resources available. We first check the conditions for extinction. Given *b*_*r*_ > *b*_*K*_ > *d*, we find that small seasonal perturbations to the resource levels cannot drive the population extinct. We next consider the conditions for fixation and loss (see Supplementary Material S1.4) and conclude that a small perturbation to resource levels in the environment cannot change the evolutionary outcome and the high-*r* allele will always be lost as in the constant environment. Numerical results show persistence of the high-*r* allele may arise under extreme changes in resource levels, however this lies outside of the scope of what is mathematically tractable in our framework.

Given these single population results, we expect local adaptation due to selection for fecundity may arise in environments with sufficiently high mortality. However, due to this high mortality and thus low population density, local adaptation may be inhibited by factors such as swamping gene flow. Therefore, we now consider when this allele can persist in a focal population with seasonal disruptions to mortality rates despite swamping gene flow and genetic drift.

### Local adaptation to seasonal population crashes

To study adaptation of a population to local seasonal disruptions, we consider an island-mainland model where the environment is constant on the mainland and seasonal on the island. Under this assumption, the mainland will consist entirely of high-*K* individuals. Migration from the mainland introduces these high-*K* alleles to the island at a constant rate *M*, hence evolution on the island can be described by the system of equations:

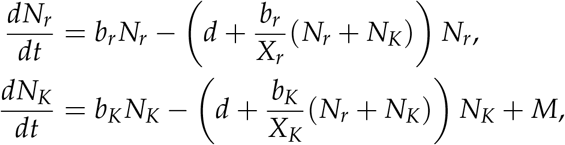

As before, we rewrite this system in terms of the frequency of the high-*r* allele on the island, *p*, and the total population size, *N*, on the island. As in Case 1 above, we incorporate seasonality by including periodic population crashes where the population density is instantaneously reduced by a proportion *δ* at the end of every season. We choose this type of seasonality for convenience, as the analysis for the case of continuous seasonality in death rates (Case 2) requires more complex mathematical techniques. As our single-population results reveal analogous conditions for *r*-selection, we expect the continuous case to yield qualitatively-similar results. These population crashes are non-selective, with individuals removed at random such that the allele frequency remains unchanged (i.e., *p*(*t*^+^) = *p*(*t*^−^), as in the single-population model). Hence, our island is modelled by the following set of differential equations with impulses:

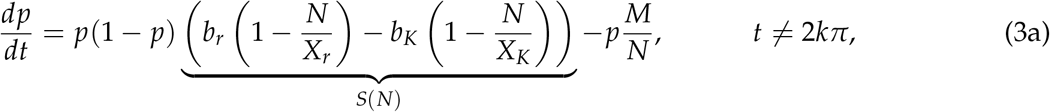

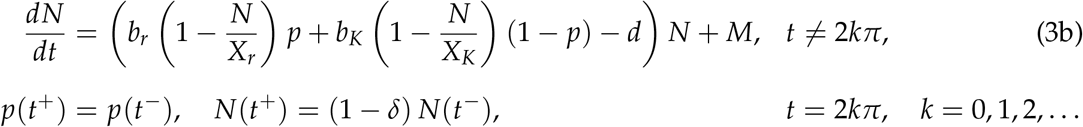

This is the same set of equations considered Case 1 of our single-population models, except for the addition of the migration term. In contrast to the single-population model, the constant influx of high-*K* alleles means that this allele can never go extinct on the island. Hence, there are only two possible long-term evolutionary outcomes. Either the high-*r* allele goes extinct so that 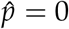, or a polymorphism persists on the island due to the balance of migration (introduction of the high-*K* allele) and selection (favouring the high-*r* allele).

#### Infinite population size limit

To determine under what conditions a polymorphism may arise, we again use Floquet analysis to consider the stability of the solution when the high-*r* allele is lost 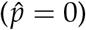 (see Supplementary Material S2.1). The persistence of the high-*r* allele is determined by the Floquet exponent:

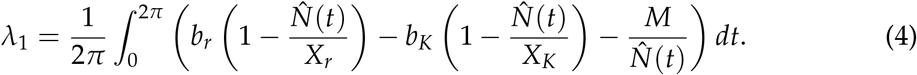

When this value is negative, the high-*r* allele will be lost. When this value is positive the high-*r* allele will persist. We calculate this value numerically across a range of parameter values (see Mathematica file for details) and compare the stability results of the island-mainland case to the minimum degree of environmental harshness, *δ*, needed for selection to favour the high-*r* allele in the single population case (Eq. 2). We call this value the *r*-selection criterion, as *δ* must be at least this large for the high-*r* allele to be favoured. Note that this criterion is not sufficient for coexistence to occur (Fig. 2A); for smaller carrying capacities, environmental harshness must be greater due to the influx of migrants which increase the overall island population size, thus requiring a harsher environment to reduce the population sufficiently to favour the high-*r* allele. We also note that if the environment is too harsh, the high-*K* individuals migrating in will outcompete the dwindling high-*r* individuals, before the environment is sufficiently harsh to drive the high-*r* alleles to extinction entirely. Thus the window in which a polymorphism persists and the island becomes locally adapted to favour the high-*r* allele may be smaller than the window in which the high-*r* allele is favoured in a single population model. As carrying capacity increases (Fig. 2B), the parameter region in which coexistence occurs grows to match the parameter region in which the high-*r* allele fixes in the single-population model. As this effect is due to the relationship between the amount of migration and the number of individuals on the island, decreasing migration will have the same effect as increasing the carrying capacity. This is in agreement with previous work showing that increasing migration decreases local adaptation (Blanquart et al., 2012).

**Figure 2:**
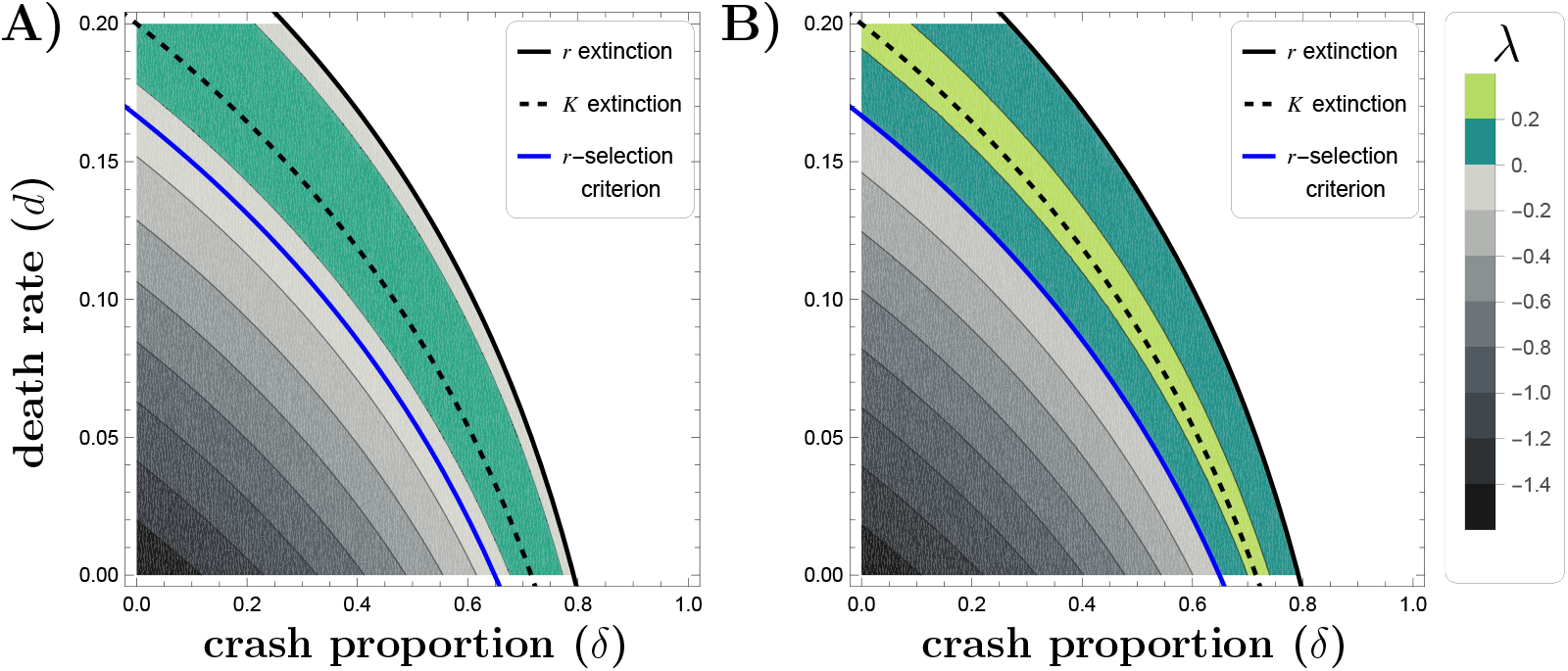
Environmental criteria for persistence of the high-*r* allele. When the Floquet exponent, *λ*, is negative (grey), the high-*r* allele is lost. When *λ* > 0 (turquoise), the high-*r* allele will persist on the island. The Floquet exponent was calculated across different degrees of environmental harshness (*δ*) and baseline death rates (*d*). Death rates ranged from 0 to *b*_*K*_. Values of *δ* were chosen below the *r* extinction criterion (black line), the maximum *δ* such that a population of high-*r* allele individuals is viable (Eq. S3). For population crashes larger than this (white), the island becomes a sink and the high-*r* allele is lost. Parameters were as follows: (A) *b*_*r*_ = 0.25, *b*_*K*_ = 0.2, *X*_*r*_ = 500, *X*_*K*_ = 1000, *M* = 1; (B) *b*_*r*_ = 0.25, *b*_*K*_ = 0.2, *X*_*r*_ = 5000, *X*_*K*_ = 10000, *M* = 1.

While the stability analysis informs when coexistence occurs, it does not describe the extent of local adaptation on the island. In the case that the high-*r* allele persists, the extent of local adaptation can be quantified using the ‘local vs. foreign’ definition of local adaptation (Blanquart et al., 2013):

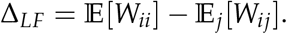

This local vs. foreign fitness contrast compares the mean fitness of individuals from the island population when reared in the island environment E[*W*_*ii*_] with the mean fitness of all individual regardless of geographic origin (e.g., *j* = *{*island, mainland*}*) when reared in the island environment (Kawecki and Ebert, 2004). In the case of the island-mainland model, this becomes:

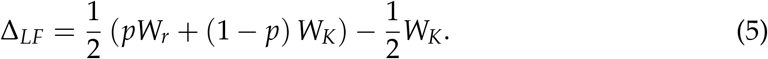

Here the fitness of an allele is given by 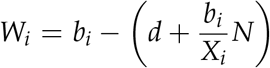.This is equivalent to the per-capita growth rate, 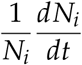. Since fitness is dependent on the population density in the environment, *N*, and allele frequency, *p*, which both change with time, it is necessary to choose a time when local adaptation will be measured. We choose to measure the degree of local adaptation on the island at both the beginning of the season and at the end, just before the crash. This will correspond to two different population densities, but the allele frequency remains the same at the beginning and end. We derive an approximation for the allele frequency in Supplementary Material S2.2.

Figure 3 shows how locally adapted the island is under different environmental conditions. At the beginning of the season, the island is always positively locally adapted, but by the end of the season the degree of local adaptation has decreased across all environments. When the island conditions are less harsh, the population may even become locally maladapted (Fig. 3). This reflects the density-dependent nature of the selection. As the season goes on, the population density increases and the high-*r* individuals become less well-suited to the environment, resulting in local maladaptation. Therefore, a population crash is necessary to maintain a low population density and hence allow the high-*r* individuals to persist.

**Figure 3:**
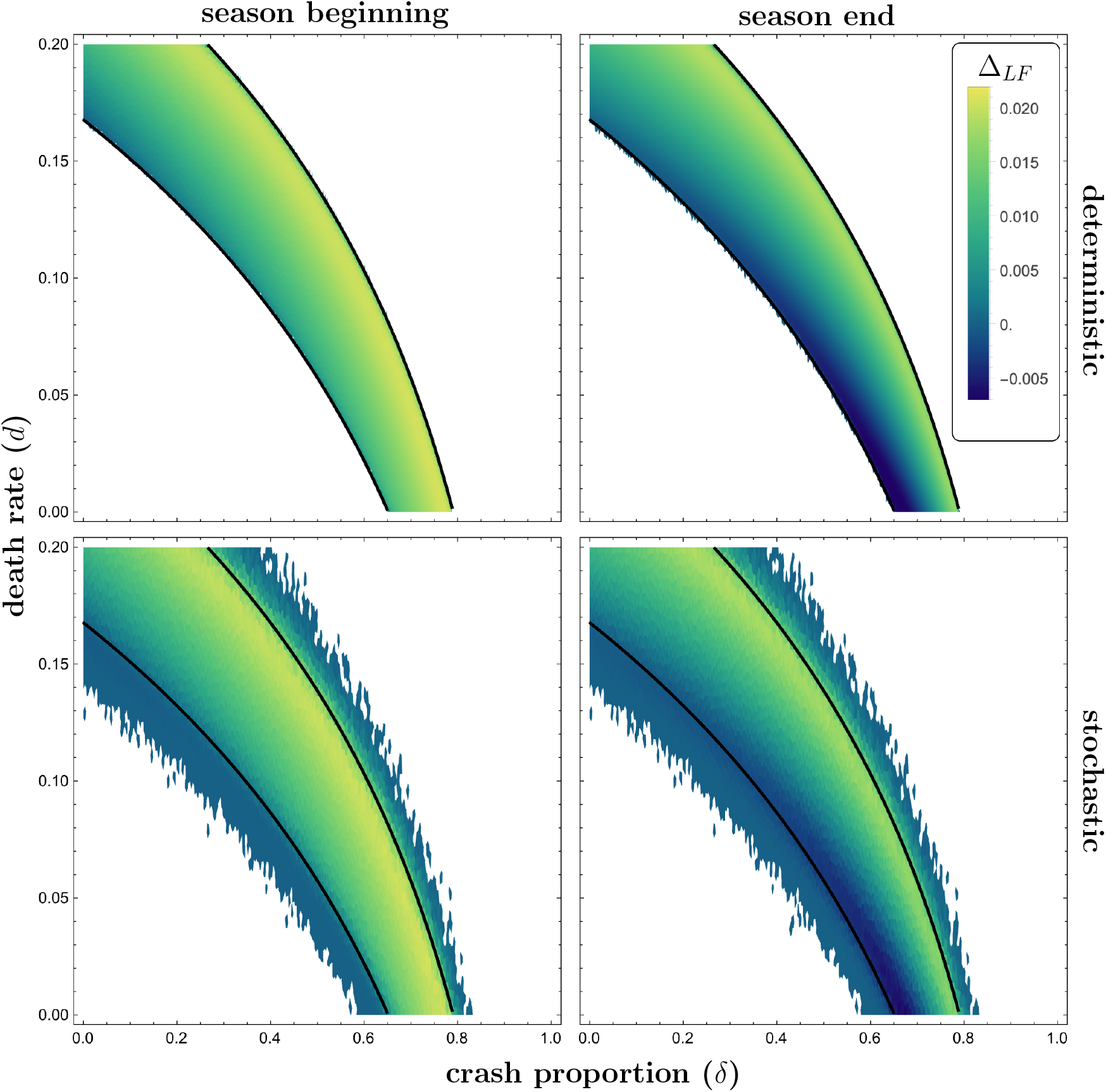
Degree of local adaptation (Δ_*LF*_) due to seasonal disruptions. Local adaptation is calculated according to Eq. 5. Black lines mark the point where the state of the high-*r* allele switches between persistence and loss, calculated numerically from the deterministic model. For both the deterministic and stochastic models, local adaptation on the island is positive at the beginning of the season. Towards the end of the season, local adaptation becomes negative in less harsh environments, indicating maladaptation. Parameters: *b*_*r*_ = 0.25, *b*_*K*_ = 0.2, *X*_*r*_ = 5000, *X*_*K*_ = 10000, *M* = 1. Stochastic simulations were initialised with *N*_*r*_ = 1000 and *N*_*K*_ = 1000, with a run time of 300.

#### Finite population size

The differential equation model presented above assumes an infinite population size, where evolution is deterministic. While this can well-approximate large populations, a real-world, finite populations are subject to stochastic effects—both evolutionary (genetic drift) and demographic—which can significantly disrupt a smaller population’s dynamics. If selection is sufficiently weak or the effective population size is sufficiently small, the evolution of a gene may be nearly neutral (Kimura, 1983). Selection is considered weak relative to neutral drift if the product of the selection coefficient *s* and the effective population size *N*_*e*_ is small, that is *sN*_*e*_ < 1 (Ohta and Tachida, 1990).

When population size fluctuates through time, as in this model, the effective population size is given by the harmonic mean over the course of a single cycle (Hartl and Clark, 2007), so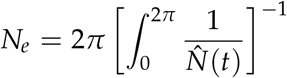. The strength of selection is measured by the selection coefficient (Eq. 3a) evaluated at the effective population size, so *s* = *S*(*N*_*e*_). We find that selection is weakest along the areas of transition between coexistence and loss of the high-*r* allele (Fig. S1). This can also be seen in the magnitude of the Floquet exponent, which is smallest along these transition regions (Fig. 2). A lower magnitude corresponds to weaker selection and thus greater susceptibility to the stochastic effects of a finite population (Yeaman and Otto, 2011). Since the environmental regime under which local adaptation arises coincides with the conditions for strong genetic drift, it is not clear that local adaptation can be maintained under finite population conditions. To understand how stochasticity might change the evolutionary outcomes in these parameter regions, we implement a finite population model of our island-mainland system.

We translate the dynamics presented in the deterministic case to a stochastic framework by implementing a Gillespie algorithm (Gillespie, 1977) with the birth, death, and migration rates found in Table 1. This allows us to capture not only the stochasticity of genetic drift, but the effects of demographic stochasticity. At the end of a season, a proportion *δ* of the population is removed. Individuals are chosen to be removed at random, with no preference for one allele over the other, hence no selection occurs. Simulations were performed across 7457 choices of the parameters *δ* and *d* with 50 replicates each.

**Table 1:** Birth, death, and migration rates.

| Event | Rate |
| --- | --- |
| birth of individual $i$ | $b_i N_i$ |
| death of individual $i$ | $\left(d + \frac{b_i}{X_i} N_T\right) N_i$ |
| migration in of high-$K$ individual | $M$ |

Note: Here $i$ refers to the genotype and $N_T$ to the total population size across all genotypes.

To quantify the local adaptation on the island, the population size and allele frequency were measured at the beginning and end of the last season of each simulation. These values were then used to calculate the local adaptation of the island at the beginning and end of the season, using the same ‘local vs. foreign’ definition (Eq. 5) as in the deterministic case. Figure 3 shows that the pattern of local adaptation on the island predicted by the deterministic result is robust to genetic drift. A transient region arises near the boundaries of transition between coexistence and loss of the high-*r* allele, in which stochasticity allows for the occasional persistence of the high-*r* allele. This results in a small, non-zero average degree of local adaptation. As the magnitude of the Floquet exponent decreases with the capacity of the island (Fig. 2A), we also predict that the effects of a finite population will increase as carrying capacity decreases.

## Discussion

Here we have used mathematical modelling and stochastic simulations to study the evolution and local adaptation of life-history traits to seasonal disruptions. Building on the results of Roughgarden (1971) in discrete time, we use a continuous-time model to show that higher fecundity can be favoured over higher competitive ability if the average amount of mortality over the course of a season reaches an intermediate level (Fig. 1C). If mortality is too low, higher competitive ability is favoured just as in a constant environment. If mortality is too high, the population will be driven to extinction. Notably, the timing of the mortality plays no role in the evolutionary outcome, with both impulsive population crashes and continuously varying death rates leading to the same outcomes. These results are in line with previous work showing *r*-selected individuals are favoured under repeated seasonal mortality (Roughgarden, 1971). Conversely, small, seasonal perturbations to the available resources in the population cannot result in selection for a high rate of reproduction. Restricting access to resources does not sufficiently reduce the population density to allow for the success of the high-growth allele. Incorporating seasonal population crashes into a model of local adaptation to an island environment relative to a mainland, we show that the conditions for local adaptation to arise resemble those necessary for *r*-selection in a single population. Simulations show that these results are robust to genetic drift and demographic stochasticity.

Fluctuations in population density due to seasonal environmental changes are already known to influence local adaptation of non-life-history traits (Pisa et al., 2019). Under periodically-changing selective forces, it has also been shown that local adaptation is maximised under intermediate levels of migration and that such periodic changes can induce the evolution of migration (Blanquart and Gandon, 2011). Theoretical work has explored local adaptation of life-history traits such as mate choice, parasite virulence, or host resistance (Best et al., 2011; Cotto et al., 2022; Gandon, 2002; Lion and Gandon, 2015), as well as the evolution of life-history traits during range expansion into a homogeneous environment (Burton et al., 2010). While the theoretical literature on local adaptation is robust, no previous work seems to exist where life-history traits, such as fecundity and competitive ability, evolve as a result of spatial variation in seasonality, although such spatial environments exist across a broad range of systems (e.g., intertidal zones, drug treatments). Favouring life-history traits requires complex environmental differences, such as changes in seasonal patterns, which poses an additional challenge in modelling studies. These environmental pressures may require strong effects on the population size, due to the density-dependent nature of these traits. Thus, it has not been clear whether the environmental conditions necessary to favour fecundity (*r*) over a tolerance for high densities (*K*) also create the conditions for the effects of drift and migration to outweigh the effects of selection, preventing local adaptation. Our results show that such a regime can exist, leading to the coexistence of *r*-selected individuals and *K*-selected individuals across space due to the balance of migration, selection, and drift.

Our theoretical findings align with observations in natural systems. Our conclusion on the effect of fluctuating resource levels may explain why the drought-intolerant strain of *L. parviflorus*, with its greater competitive ability, is dominant in non-serpentine soil, despite temporal variability in rainfall (Dittmar and Schemske, 2023). At first glance, our results appear to contradict previous theoretical work that shows *r*-selection can arise under fluctuating resource conditions (Grover, 1991; Hsu, 1980; Kremer and Klausmeier, 2013; Litchman and Klausmeier, 2001; Tachikawa, 2008). However, in all these models changes to resource levels resulted in large and immediate fluctuations in the birth rate. This has a direct impact on the population size, causing large decreases in the population density which allows for *r*-selected individuals to out-compete *K*-selected individuals as the population reestablishes itself. By contrast, our model assumes a small perturbation to resources that results in gradual, linearly-dependent changes in density-dependent mortality. In this case, changes in population size are insufficient to favour the *r*-selected allele. Therefore, we conclude that the effects of seasonal disruptions to resource levels are dependent on the speed and severity of the responding fluctuations in population demography.

The seaweed fly *Coelopa frigida* is an excellent case study for *r*and *K*-selection. It displays three phenotypes: a homozygote with a large body size and slow maturation, a homozygote with a small body size and fast maturation, and an intermediate heterozygote. *C. frigida* is a noteworthy case in part because of a chromosomal inversion that allows the group of genes that confer these *r*- or *K*-selected phenotypes to be passed onto offspring together (Mérot et al., 2018, 2020). Thus the inheritance pattern functionally behaves like that of a single biallelic locus, as in the model presented here. However such patterns of inheritance are rare; more often *r*- and *K*-selection emerges from a combination of phenotypes, all mediated by multiple genes. The simplifying assumption of a single locus model allows for better tractability and focuses on the question of what environmental conditions can or cannot lead to broad patterns of *r*-selection over *K*-selection. Determining how more complex genetic architecture or trade-offs between more specific phenotypic traits may complicate the relationship of *r*- and *K*-selection and seasonality requires more complicated models and empirical work. Notably, a model which allows for a greater variety of phenotypes would show how intermediate life-history strategies, which are neither entirely *r*-selected nor entirely *K*-selected, might respond to different environmental disruptions.

The models presented here describe the evolution of only one of many possible lifehistory trade-offs shaped by density-dependent natural selection. Specifically, the models considered did not incorporate the age structure that would be necessary for modelling to fully capture life-history evolution, as seasonality and mortality frequently both relate to the life cycle and age of individuals. A model incorporating age structure would explore questions of selective pressures on the rate of maturation and the importance of the timing of mortality relative to the life cycle. Additionally, these models assumed regular seasonality with a consistent severity. However, a consequence of climate change is irregular seasonality and more dramatic seasonal swings. As the work of Lande et al. has demonstrated, stochasticity also serves to shape life-history evolutionary dynamics (Lande et al., 2009, 2017). Incorporating a degree of stochasticity into the timing and severity of seasons may change the dynamics shown here. Alternatively, some environments experience seasons of great stability and seasons of large disruptions, motivating a model consisting of seasons of environmental stability and seasons of regular disruption. Both these cases may be of particular interest for stochastic simulations, to study the transient states in regions of higher or lower mortality (Fig. 3) where dynamics may easily shift in response to greater variability.

Given our focus on a haploid single-locus life-history trait, our examination of lifehistory evolution within a single population is closely related to the study of species coexistence in variable environments (Chesson, 2000). This comparison highlights a key assumption of our model: the Lotka-Volterra model used here assumes that the death rate increases linearly with population density (a proxy for declining resource availability). Such linear (e.g., Type I functional responses) dependence on population size can never result in stable polymorphism (e.g., coexistence of the *r*-selected and *K*-selected species). Results from the coexistence theory suggest that introducing a non-linear (e.g., Type II) response can facilitate coexistence (Armstrong and McGehee, 1980). While our single-population results suggest that linear responses do not facilitate coexistence within isolated patches, the examination of local adaptation indicates that these fluctuations shape when species can appear to coexist as a result of continued immigration. Nevertheless, building on the findings of coexistence theory, additional work is needed to understand when local adaptation can occur when there are non-linear responses to available resources or when life-history traits have more complex multi-locus genetic bases whose dynamics may not be well predicted by species coexistence.

The work presented here extends classic life-history theory (Anderson, 1971; MacArthur and Wilson, 1967; Roughgarden, 1971), demonstrating how the seasonality can (in the case of seasonal mortality) and cannot (in the case of fluctuating resource abundance) favour the spread of *r*-selected life-history strategies. We show that *r*-selected life-history traits are not favoured in all seasonal environments, but rather selection depends on the nature (i.e., how seasonality impacts populations by changing population size, resource availability, or mortality rate), timing, and severity of seasonal disruptions. These results can be useful for understanding life-history evolution in experimental, agricultural, and natural systems. As anthropogenic climate change has increasingly dramatic effects on seasonality and variability, understanding how temporal environmental variation shapes evolution in single populations as well as local adaptation across spatially-structured environments is increasingly important. The results and methods presented here establish a foundation on which this future work can build, to further elucidate the complex ecoevolutionary consequences of seasonality on the evolution of density-dependent traits.

## Supporting information

Mathematica Notebook PDF

## Acknowledgements

RH and AM were supported by funds from the National Science and Engineering Research Council of Canada [NSERC: RGPIN-2022-03113, CRC-2021-00276]. In addition RH was supported by the SFU Entrance Scholarship and Peter Borwein Memorial Scholarship from Simon Fraser University. The authors would like to thank Ben Ashby, Leithen M’Gonigle, Puneeth Deraje, Kuangyi Xu, Chris Carlson, and Mete Yuksel for their helpful feedback on the manuscript.

