## Supplemental Material for "Local Adaptation of Life-History Traits in a Seasonal Environment"

### Supplementary Figures

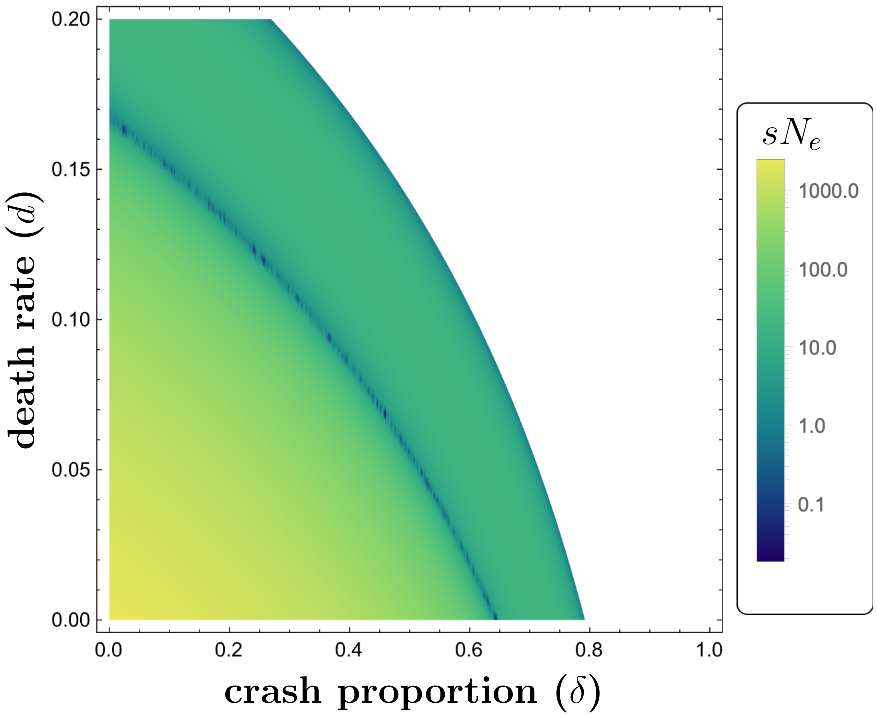


Figure S1: **Product of the selection coefficient and the effective population size.** For values near or less than 1, selection is considered weak relative to neutral drift. $sN_e$ was not calculated in the parameter region where the island is a sink (white). Note values are shown on a log scale. Parameters: $b_r = 0.25$, $b_K = 0.2$, $X_r = 5000$, $X_K = 10000$, $M = 1$.

### S1: Single population model

We begin by considering the evolution of $r$ vs. $K$ life-history traits in a single population. We model population growth with a logistic model where the population size $N$ is given by the differential equation:

$$\frac{dN}{dt} = bN - \left(d + \frac{b}{X}N\right)N,$$

where $b$ is the per-capita birth rate, $d$ is the death rate, and $X$ is the 'growth limit' which determines the carrying capacity (a.k.a. equilibrium) population size $K$ in the long-term:

$$K = \frac{(b-d)X}{b},$$

We additionally note that the intrinsic growth rate of the population is $r = b - d$. To model evolution in this population, we consider the dynamics at a single biallelic, haploid locus with a high-$r$ allele and a high-$K$ allele. We choose parameters such that the high-$r$ allele confers a lower carrying capacity ($K_r < K_K$), but a higher birth rate ($b_r > b_K$). The death rate ($d$) remains constant across both alleles and must be smaller than both birth rates. Hence we have:

$$0 < d < b_K < b_r,$$
$$\frac{(b_r-d)X_r}{b_r} = K_r < K_K = \frac{(b_K-d)X_K}{b_K}.$$

Evolution is described by a Lotka-Volterra model of competition between individuals carrying the high-$r$ allele and individuals carrying the high-$K$ allele, as given by the following system of differential equations:

$$\frac{dN_r}{dt} = b_r N_r - \left(d + \frac{b_r}{X_r}(N_r + N_K)\right) N_r,$$
$$\frac{dN_K}{dt} = b_K N_K - \left(d + \frac{b_K}{X_K}(N_r + N_K)\right) N_K,$$

where $N_r$ is the number of individuals carrying the high-$r$ allele and $N_K$ is the number with the high-$K$ allele.

It is convenient to re-parametrise these two equations in terms of the frequency of the high-$r$ allele, $p$, and the total population size, $N$:

$$p = \frac{N_r}{N_r + N_K},$$
$$N = N_r + N_K.$$

Hence changes in the allele frequency and changes in the total population size can be described by the following differential equations:

$$\frac{dp}{dt} = p(1-p)\left(b_r\left(1 - \frac{N}{X_r}\right) - b_K\left(1 - \frac{N}{X_K}\right)\right) = f_1(p,N) \tag{S1a}$$
$$\frac{dN}{dt} = \left(b_r\left(1 - \frac{N}{X_r}\right)p + b_K\left(1 - \frac{N}{X_K}\right)(1-p) - d\right)N = f_2(p,N). \tag{S1b}$$

Note that Eq. S1a can also be written as,

$$\frac{dp}{dt} = p(1-p)S(N),$$

where $S(N)$ is the selection coefficient given by:

$$S(N) = b_r\left(1 - \frac{N}{X_r}\right) - b_K\left(1 - \frac{N}{X_K}\right). \tag{S2}$$

System S1 exhibits three equilibria:

$$\begin{aligned}
\hat{N} &= 0 && \text{(extinction)}\\
(\hat{p}, \hat{N}) &= \left(1, \frac{(b_r - d)X_r}{b_r}\right) && \text{(fixation)}\\
(\hat{p}, \hat{N}) &= \left(0, \frac{(b_K - d)X_K}{b_K}\right) && \text{(loss).}
\end{aligned}$$

Extinction is unstable when $b_i > d$ (a condition which is assumed to be met throughout). Fixation of the high-$r$ allele is always unstable, while its loss is always stable. This agrees with classic theory which holds that in a constant environment, the high-$K$ allele will out-compete the high-$r$ allele and drive it to extinction.

### *S1.1: Seasonal population crashes: an impulsive differential equation approach*

In order to incorporate seasonal disruptions, we first consider a case where the population undergoes sudden, periodic population crashes. At the end of every season, a proportion $\delta$ of the population is removed. The length of the season is arbitrary, as the birth and death rates can be scaled. To maintain consistency across all our models, we choose a season length of $2\pi$. This model can be described by modifying System S1 to get the following set of impulsive differential equations:

$$\begin{aligned}
\frac{dp}{dt} &= f_1(p, N), && t \neq 2k\pi,\\
\frac{dN}{dt} &= f_2(p, N), && t \neq 2k\pi, \quad k = 0, 1, 2, \ldots,\\
p(t) = p(t^-), \quad N(t) &= (1-\delta)\,N(t^-), && t = 2k\pi,
\end{aligned}$$

where $p(t^-)$ and $N(t^-)$ are the allele frequency and the population size immediately before the crash. Note that $p(t^-) = p(t)$, so that the population crash does not change the allele frequency in the population. Hence, there is no selective element to the disruptions. By Theorem 5.1 from Liu et al. (2007), there is no positive periodic solution with coexistence. There are therefore three possible long-term outcomes: extinction, fixation, or loss. By Lemma 2.1 from Liu et al. (2007), if:

$$b_K - d < b_r - d < -\frac{1}{2\pi}\ln(1-\delta), \tag{S3}$$

then the population will go extinct. Otherwise, there are two semi-trivial positive periodic solutions, $\left(0, \hat{N}_K\right)$ and $\left(1, \hat{N}_r\right)$, where where $\hat{N}_i(t)$ is the positive periodic solution to:

$$\begin{aligned} \frac{dN_i}{dt} &= b_i N_i - \left(d + \frac{b_i}{X_i}N\right) N_i, && t \neq 2k\pi, \\ N_i(t) &= (1-\delta)\, N_i(t^-), && t = 2k\pi. \end{aligned}$$

To find the linear stability of each of these solutions we must compute the mean population over a season:

$$\mathbb{E}(\hat{N}_i) = \frac{1}{2\pi}\int_0^{2\pi} \hat{N}_i(t)dt = \frac{X_i}{b_i}\left((b_i - d) + \frac{1}{2\pi}\ln(1-\delta)\right).$$

For the sake of clean notation, we use expectation notation rather than the bar notation generally used to denote the mean. Using Theorem 4.2 from Liu et al. (2007), we can see that the $\hat{p} = 0$ case will be linearly stable if:

$$\begin{aligned} b_K - d &> -\frac{1}{2\pi}\ln(1-\delta) \\ b_r - d &< -\frac{1}{2\pi}\ln(1-\delta) + \frac{b_r}{X_r}\mathbb{E}(\hat{N}_i). \end{aligned}$$

Similarly, the $\hat{p} = 1$ case will be linearly stable if:

$$\begin{aligned} b_r - d &> -\frac{1}{2\pi}\ln(1-\delta) \\ b_K - d &< -\frac{1}{2\pi}\ln(1-\delta) + \frac{b_K}{X_K}\mathbb{E}(\hat{N}_i). \end{aligned}$$

By plugging $\mathbb{E}(\hat{N}_i)$ into our stability condition and rearranging, we find that the $\hat{p} = 0$ case is stable if:

$$\delta < 1 - \exp\left[-2\pi(b_K - d)\right]$$
$$\delta < 1 - \exp\left[-2\pi\left(\frac{b_r X_K(b_K - d) - b_K X_r(b_r - d)}{b_r X_K - b_K X_r}\right)\right].$$

The equilibrium solution $\hat{p} = 1$ is stable if:

$$\delta < 1 - \exp\left[-2\pi(b_r - d)\right]$$
$$\delta > 1 - \exp\left[-2\pi\left(\frac{b_r X_K(b_K - d) - b_K X_r(b_r - d)}{b_r X_K - b_K X_r}\right)\right].$$

Given the restrictions on our parameters, we have that the above bounds are restricted to the order:

$$1 - \exp\left[-2\pi\left(\frac{b_r X_K(b_K - d) - b_K X_r(b_r - d)}{b_r X_K - b_K X_r}\right)\right] < 1 - \exp\left[-2\pi\left(b_K - d\right)\right] < 1 - \exp\left[-2\pi\left(b_r - d\right)\right].$$

Hence we find that the $\hat{p} = 1$ case, when the high-$r$ allele overtakes the population, is linearly stable in the range:

$$1 - \exp\left[-2\pi\left(\frac{b_r X_K(b_K - d) - b_K X_r(b_r - d)}{b_r X_K - b_K X_r}\right)\right] < \delta < 1 - \exp\left[-2\pi(b_r - d)\right]. \quad \text{(S4)}$$

If the magnitude of the population crashes is smaller, then the high-$K$ allele will overtake the population (i.e., $\hat{p} = 0$ is stable). If the population crashes are too large, the population will go extinct.

### *S1.2: Floquet analysis for continuous disruptions*

In the following two sections, we will consider continuous seasonal changes. The population dynamics under these conditions can be described by a small, periodic perturbation to the original system described by System S1:

$$\frac{dp}{dt} = f_1(p, N) - \epsilon g_1(p, N)$$
$$\frac{dN}{dt} = f_2(p, N) - \epsilon g_2(p, N),$$

where $\epsilon$ is a small, positive perturbation and $g_i(p, N)$ are periodic functions. By incorporating any periodic perturbation, the system will no longer reach a constant equilibrium. Rather, we must consider the periodic equilibrium solutions and their stability. To do so, we apply Floquet analysis (Klausmeier, 2008).

Let $(\hat{p}, \hat{N})$ be a periodic solution. Then we consider the behaviour of a small perturbation to this solution: $(p(t), N(t)) = (\hat{p} + \xi_1(t), \hat{N} + \xi_2(t))$. These perturbations may be written as:

$$\begin{pmatrix} \xi_1(t) \\ \xi_2(t) \end{pmatrix} = \Phi(t) \begin{pmatrix} \xi_1(0) \\ \xi_2(0) \end{pmatrix},$$

where $\Phi(t)$ is the matrix satisfying:

$$\frac{d\Phi}{dt} = \mathbf{A}\big|_{p=\hat{p}, N=\hat{N}} \Phi(t), \quad \Phi(0) = \mathbf{I},$$

and $\mathbf{A}$ is the matrix:

$$\mathbf{A} = \begin{pmatrix} \frac{\partial}{\partial p}\left(f_1(p, N) - \epsilon g_1(p, N)\right) & \frac{\partial}{\partial N}\left(f_1(p, N) - \epsilon g_1(p, N)\right) \\ \frac{\partial}{\partial p}\left(f_2(p, N) - \epsilon g_2(p, N)\right) & \frac{\partial}{\partial N}\left(f_2(p, N) - \epsilon g_2(p, N)\right) \end{pmatrix}.$$

Then the Floquet multipliers, $\rho_i$, of the system are the eigenvalues of $\Phi(2\pi)$. The periodic solution $(\hat{p}, \hat{N})$ is locally stable if both of its Floquet multipliers satisfy:

$$|\rho_i| < 1.$$

In some cases, it may be easier to look at the Floquet exponents, $\lambda_i$, which are derived from the Floquet multipliers by:

$$\lambda_i = \frac{1}{2\pi} \ln |\rho_i|.$$

In order for the periodic solution to be locally stable, these Floquet exponents must be negative.

### *S1.3: Continuous seasonal variations in death rates*

To model continuous variation in the death rate, we substitute $d \mapsto d(1 + \epsilon h(t))$, where $h(t)$ is a periodic function and $\epsilon \in (0, 1)$ is a small quantity. This restriction of $\epsilon$ indicates that we are considering a death rate that depends weakly on time and ensures that the death rate is always positive. Without loss of generality, we may assume that $h(t)$ has a period of $2\pi$. For longer or shorter seasons, the birth and death rates can be scaled. This results in the perturbed problem:

$$\frac{dp}{dt} = f_1(p, N)$$
$$\frac{dN}{dt} = f_2(p, N) - \epsilon d N h(t).$$

We could consider a specific periodic function $h(t)$, but as we will show below, the result is sensitive to the shape of this function. Hence we take a general approach where we express $h(t)$ as a Fourier series:

$$h(t) = A_0 + \sum_{n=1}^{\infty} A_n \cos nt + B_n \sin nt,$$

where:

$$A_0 = \frac{1}{2\pi} \int_0^{2\pi} h(t) dt,$$
$$A_n = \frac{1}{\pi} \int_0^{2\pi} h(t) \cos(nt) dt,$$
$$B_n = \frac{1}{\pi} \int_0^{2\pi} h(t) \sin(nt) dt.$$

The Floquet exponents for this system are:

$$\lambda_1 = \frac{1}{2\pi} \int_0^{2\pi} \frac{\partial}{\partial N} f_2(p(t), N(t)) - \epsilon d h(t) dt \quad \text{(S5)}$$
$$\lambda_2 = \frac{1}{2\pi} \int_0^{2\pi} (1 - 2p(t))\, S\left(N(t)\right) dt, \quad \text{(S6)}$$

where $S(N)$ is the selection coefficient given in Eq. S2. We seek that Eqs. S5 and S6 be negative to ensure the local stability of the solution.

We first check the stability conditions for the equilibrium states when the population goes extinct, $\left(\hat{p}, \hat{N}\right) = (0, 0)$ and $\left(\hat{p}, \hat{N}\right) = (1, 0)$. Given $b_r > b_K$, we find Eq. S6 is always positive for $\left(\hat{p}, \hat{N}\right) = (0, 0)$, so this state is always unstable. For $\left(\hat{p}, \hat{N}\right) = (1, 0)$, Eq. S6 is always negative. From Eq. S5 we find that the condition for $\left(\hat{p}, \hat{N}\right) = (1, 0)$ to be stable is:

$$\epsilon A_0 > \frac{b_r}{d} - 1.$$

This is our condition for extinction.

To determine which genotype will dominate when the population is not driven to extinction, we first consider the equilibrium at $\hat{p} = 0$, when the high-$r$ allele is lost. Recall that in the unperturbed problem, this is always the stable steady state. To consider the periodic equilibrium solution for the population size in this case, we take the asymptotic expansion:

$$\hat{N} = N_0 + \epsilon N_1 + \mathcal{O}(\epsilon^2),$$

and plug this into the perturbed equation for population size:

$$\frac{dN}{dt} = \left(b_K \left(1 - \frac{N}{X_K}\right) - d\right) N - \epsilon d h(t) N.$$

At $\mathcal{O}(1)$, we arrive at the unperturbed problem. We take the leading order of our expansion to be the stable steady state, so:

$$N_0 = \frac{(b_K - d)X_K}{b_K}.$$

At $\mathcal{O}(\epsilon)$, we get the differential equation:

$$\frac{dN_1}{dt} = -(b_K - d)N_1 - dN_0 \left[A_0 + \sum_{n=1}^{\infty} A_n \cos nt + B_n \sin nt\right]. \tag{S7}$$

Since we are looking for a periodic solution, we assume that $N_1$ can be written as a Fourier series:

$$N_1(t) = a_0 + \sum_{n=1}^{\infty} a_n \cos nt + b_n \sin nt.$$

Plugging this into Eq. S7 and taking $\alpha = -(b_K - d)$ and $\beta = dN_0$, we find that:

$$\alpha a_0 - \beta A_0 + \sum_{n=1}^{\infty} (\alpha a_n - \beta A_n - n b_n) \cos nt + (\alpha b_n - \beta B_n + n a_n) \sin nt = 0.$$

This implies that:

$$\alpha a_0 - \beta A_0 = 0,$$

$$\alpha a_n - \beta A_n - n b_a = 0,$$

$$\alpha b_n - \beta B_n + n a_n = 0,$$

hence:

$$
\begin{aligned}
a_0 &= \frac{\beta}{\alpha} A_0, \\
a_n &= \frac{\beta\left(\alpha A_n + n B_n\right)}{\alpha^2 + n^2}, \\
b_n &= \frac{\beta\left(\alpha B_n - n A_n\right)}{\alpha^2 + n^2}.
\end{aligned}
$$

Therefore:

$$N_1 = \frac{\beta}{\alpha} A_0 + \sum_{n=1}^{\infty} \frac{\beta\left(\alpha A_n + n B_n\right)}{\alpha^2 + n^2} \cos nt + \frac{\beta\left(\alpha B_n - n A_n\right)}{\alpha^2 + n^2} \sin nt,$$

which gives us the asymptotic expansion of $\hat{N}$ near the steady state of the unperturbed problem:

$$\hat{N} = N_0 + \epsilon \left[ \frac{\beta}{\alpha} A_0 + \sum_{n=1}^{\infty} \frac{\beta\left(\alpha A_n + n B_n\right)}{\alpha^2 + n^2} \cos nt + \frac{\beta\left(\alpha B_n - n A_n\right)}{\alpha^2 + n^2} \sin nt \right] + \mathcal{O}(\epsilon^2).$$

Plugging $\hat{p} = 0$ into our first Floquet exponent (Eq. S5), we have:

$$\lambda_1 = \frac{1}{2\pi} \int_0^{2\pi} b_K - d - \frac{2b_K}{X_K} \hat{N}(t) - \epsilon d h(t) dt.$$

This requires us to compute:

$$
\begin{aligned}
\int_0^{2\pi} \hat{N} &= \int_0^{2\pi} N_0 + \epsilon \left[ \frac{\beta}{\alpha} A_0 + \sum_{n=1}^{\infty} a_n \cos nt + b_n \sin nt \right] dt \\
&= 2\pi \left( N_0 + \epsilon \frac{\beta}{\alpha} A_0 \right) + \epsilon \sum_{n=1}^{\infty} \left[ a_n \int_0^{2\pi} \cos(nt) dt + b_n \int_0^{2\pi} \sin(nt) dt \right] \\
&= 2\pi \left( N_0 + \epsilon \frac{\beta}{\alpha} A_0 \right).
\end{aligned}
\tag{S8}
$$

Plugging Eq. S8 into our Floquet exponent and recalling that $A_0 = \frac{1}{2\pi} \int_0^{2\pi} h(t) dt$, we find that:

$$\lambda_1 = -\left[ b_K - d(1 + \epsilon A_0) \right].$$

This is negative so long as:

$$\epsilon A_0 < \frac{b_K}{d} - 1.$$

It remains to check whether the direction of selection will change due to the seasonal perturbation. We therefore check the second Floquet exponent (Eq. S6). Substituting $\alpha = -(b_K - d)$

and $\beta = dN_0$ back in and using the result in Eq. S8, we find that the second exponent is negative when the following inequality holds:

$$\epsilon A_0 < \frac{b_r X_K (b_K - d) - b_K X_r (b_r - d)}{d(b_r X_K - b_K X_r)}. \tag{S9}$$

Since:

$$\frac{b_r X_K (b_K - d) - b_K X_r (b_r - d)}{d(b_r X_K - b_K X_r)} < \frac{b_K}{d} - 1,$$

we see that $\hat{p} = 0$ is stable whenever Eq. S9 holds.

To consider the stability of $\hat{p} = 1$, we follow the same steps and find that the first Floquet exponent (Eq. S5) is negative when:

$$\epsilon A_0 < \frac{b_r}{d} - 1,$$

and the second Floquet exponent (Eq. S6) is negative when Eq. S9 does not hold. Since:

$$\frac{b_r X_K (b_K - d) - b_K X_r (b_r - d)}{d(b_r X_K - b_K X_r)} < \frac{b_r}{d} - 1,$$

the $\hat{p} = 1$ solution is stable when $\epsilon A_0$ is between these two values. Thus, the high-$r$ allele will dominate the population when:

$$\frac{b_r X_K (b_K - d) - b_K X_r (b_r - d)}{d(b_r X_K - b_K X_r)} < \epsilon A_0 < \frac{b_r}{d} - 1. \tag{S10}$$

In this case, the seasonal perturbations to the death rate are large enough and long enough to favour high-$r$ strategies, but not so harsh that the population is driven extinct. Note that as the baseline death rate, $d$, increases, this window for $\epsilon A_0$ shifts so that seasonal perturbations need not be as harsh. We also note that Eq. S10 can equivalently be written as:

$$1 - \exp\left[-2\pi \left(\frac{b_r X_K (b_K - d) - b_K X_r (b_r - d)}{b_r X_K - b_K X_r}\right)\right] < 1 - e^{-2\pi d \epsilon A_0} < 1 - \exp\left[-2\pi (b_r - d)\right],$$

which is the same inequality as Eq. S4, except with $\delta$ replaced by $1 - e^{-2\pi d \epsilon A_0}$. This is the probability of there being a death in $2\pi$ time in a Poisson process with a death rate of $d\epsilon A_0$. Given that $d\epsilon A_0$ is the averaged size of the perturbation to the death rate, this can be interpreted as the proportion of the population killed due to the perturbed death rate. Hence $\delta$ and $1 - e^{-2\pi d \epsilon A_0}$ both represent the proportion of the population that dies over the course of the season above what would be expected from the natural death rate.

### *S1.4: Continuous seasonal variations in resource levels*

To model continuous variation in the resource levels, we apply a periodic change to the carrying capacity by substituting $X_i \mapsto X_i(1-\epsilon h(t))$, where $h(t)$ is a periodic function. Without loss of generality, we again assume that $h(t)$ has a period of $2\pi$. In this case, we have a perturbation in the denominator. That is, $\dfrac{1}{X_i}$ becomes:

$$\frac{1}{X_i(1-\epsilon h(t))}.$$

Therefore, in order to carry on with our perturbation analysis as before, we must first perform a Taylor expansion so that:

$$\frac{1}{X_i(1-\epsilon h(t))} \approx X_i^{-1}(1+\epsilon h(t)).$$

Substituting in this expansion and rewriting our functions, we arrive at the perturbed problem:

$$\begin{aligned}\frac{dp}{dt} &= f_1(p,N) - \epsilon p(1-p)\left(\frac{b_r}{X_r} - \frac{b_K}{X_K}\right) N h(t)\\ \frac{dN}{dt} &= f_2(p,N) - \epsilon\left[p\frac{b_r}{X_r} - (1-p)\frac{b_K}{X_K}\right] N^2 h(t).\end{aligned}$$

We express $h(t)$ as a Fourier series as before. Recall that from this we have:

$$A_0 = \frac{1}{2\pi}\int_0^{2\pi} h(t)dt.$$

Now the Floquet exponents for this system are:

$$\lambda_1 = \frac{1}{2\pi}\int_0^{2\pi} \frac{\partial}{\partial N} f_2(p(t),N(t)) - \epsilon\left(2\frac{b_r}{X_r}p(t) - 2\frac{b_K}{X_K}(1-p(t))\right) N(t)h(t)dt, \tag{S11}$$

$$\lambda_2 = \frac{1}{2\pi}\int_0^{2\pi} (1-2p(t))S(N(t)) - \epsilon(1-2p(t))\left(\frac{b_r}{X_r} - \frac{b_K}{X_K}\right) N(t)h(t)dt. \tag{S12}$$

These must be negative to ensure local stability.

We first check the extinction cases: $(\hat{p},\hat{N}) = (0,0)$ and $(\hat{p},\hat{N}) = (1,0)$. Given $b_r > b_K > d$, Eq. S11 is always positive for both extinction cases. Therefore, seasonal perturbations to the resource levels cannot drive the population extinct.

We next consider the equilibrium at $\hat{p} = 0$ when the high-$r$ allele is lost. In the unperturbed problem, this is always the stable steady state. Once again, we take the asymptotic expansion of $\hat{N}(t)$ and plug this into the perturbed equation for population size:

$$\frac{dN}{dt} = \left(b_K\left(1 - \frac{N}{X_K}\right) - d\right)N - \epsilon\frac{b_K}{X_K}h(t)N.$$

At $\mathcal{O}(1)$, we have the unperturbed problem and take the leading order of our expansion to be its stable steady state. At $\mathcal{O}(\epsilon)$, we get the differential equation:

$$\frac{dN_1}{dt} = \left(b_K - d - 2\frac{b_K}{X_K}N_0\right)N_1 - \frac{b_K}{X_K}N_0\left[A_0 + \sum_{n=1}^{\infty} A_n \cos nt + B_n \sin nt\right]. \tag{S13}$$

Since we are looking for a periodic solution, we again require that $N_1$ can be written as a Fourier series:

$$N_1(t) = a_0 + \sum_{n=1}^{\infty} a_n \cos nt + b_n \sin nt.$$

Plugging this into Eq. S13 and taking $\alpha = -(b_K - d)$ and $\beta = (b_K - d)N_0$, we find that:

$$\alpha a_0 - \beta A_0 + \sum_{n=1}^{\infty}(\alpha a_n - \beta A_n - n b_n)\cos nt + (\alpha b_n - \beta B_n + n a_n)\sin nt = 0.$$

This implies that:

$$\begin{aligned}
\alpha a_0 - \beta A_0 &= 0,\\
\alpha a_n - \beta A_n - n b_n &= 0,\\
\alpha b_n - \beta B_n + n a_n &= 0,
\end{aligned}$$

hence:

$$\begin{aligned}
a_0 &= \frac{\beta}{\alpha}A_0,\\
a_n &= \frac{\beta(\alpha A_n + n B_n)}{\alpha^2 + n^2},\\
b_n &= \frac{\beta(\alpha B_n - n A_n)}{\alpha^2 + n^2}.
\end{aligned}$$

Altogether, we have the asymptotic expansion of $\hat{N}$ near the steady state of the unperturbed problem:

$$\hat{N} = N_0 + \epsilon\left[\frac{\beta}{\alpha}A_0 + \sum_{n=1}^{\infty}\frac{\beta(\alpha A_n + n B_n)}{\alpha^2 + n^2}\cos nt + \frac{\beta(\alpha B_n - n A_n)}{\alpha^2 + n^2}\sin nt\right] + \mathcal{O}(\epsilon^2).$$

The integral of this is the same as in Eq. S8 from the previous case.

We now plug $\hat{p} = 0$ and $\hat{N}$ into our first Floquet exponent (Eq. S11). Dropping all terms that are $\mathcal{O}(\epsilon^2)$ or smaller and using Eq. S8, we get:

$$\begin{aligned} \lambda_1 &= \frac{1}{2\pi} \int_0^{2\pi} b_K - d - 2\frac{b_K}{X_K}\hat{N} + \epsilon \left( 2\frac{b_K}{X_K} N_0 h(t) \right) dt \\ &= -\left(b_K - d\right), \end{aligned}$$

which is always negative as we have already required that $d < b_K$.

Similarly for the second Floquet exponent (Eq. S12), we find that:

$$\begin{aligned} \lambda_2 &= \frac{1}{2\pi} \int_0^{2\pi} b_r - b_K \left( \frac{b_r}{X_r} - \frac{b_K}{X_K} \right) \hat{N} - \epsilon \left( \frac{b_r}{X_r} - \frac{b_K}{X_K} \right) N_0 h(t) dt \\ &= \left( b_r - b_K \left( \frac{b_r}{X_r} \right) N_0 \right) \\ &= \left( S(N_0) \right), \end{aligned}$$

where $S(N)$ is the selection coefficient in the unperturbed environment (Eq. S2). Since we have chosen our parameters to favour the high-$K$ allele in the unperturbed environment, we know this value to be negative. Thus we find that the equilibrium at $\hat{p} = 0$ is locally stable regardless of any perturbation to the environment. Carrying out a similar analysis for the $\hat{p} = 1$ case, we find that this case is always locally unstable. Therefore, we conclude that a perturbation to resource levels in the environment cannot change the evolutionary outcome.

### S2: Island-mainland model

#### *S2.1: Floquet analysis*

We again apply Floquet analysis (Klausmeier, 2008) to study the stability of the periodic solution when $\hat{p} = 0$. In the long-term, this periodic solution must consist of repeated cycles where the population size grows logistically from an initial size to some maximum immediately before the crash. To obtain an identical cycle in the next time interval, the population must crash down to the same initial density as at the beginning of the cycle. Hence we can solve for this periodic solution by solving for the general solution from time $0 \leq t \leq 2\pi$ and then solving for the initial condition $N_0$ that results in the required limit cycle. In this case, $\hat{N}(t)$ is the solution to:

$$\frac{dN}{dt} = b_K N - \left(d + \frac{N}{X_K}\right) N + M, \quad 0 \leq t \leq 2\pi, \quad N(0) = (1-\delta)N(2\pi),$$

which is given by:

$$\hat{N}(t) = \frac{C_2 N_0 \left(1 + e^{C_1 t}\right) - \left(2M + (b_K - d)N_0\right) X_K \left(1 - e^{C_1 t}\right)}{C_2 \left(1 + e^{C_1 t}\right) - \left(2b_K N_0 - X_K(b_K - d)\right)\left(1 - e^{C_1 t}\right)},$$

where:

$$C_1 = \frac{\sqrt{4b_K M + (b_K - d)^2 X_K}}{\sqrt{X_K}},$$
$$C_2 = \sqrt{X_K}\sqrt{4b_K M + (b_K - d)^2 X_K},$$

and $N_0$ is such that:

$$N_0 = \hat{N}(0) = (1-\delta)\hat{N}(2\pi),$$

that is:

$$N_0 = \frac{X_K}{2}\left(1 - \frac{d}{b_K}\right)\left(1 - \frac{\delta}{2}\right) - \frac{(e^{C_1\pi} + e^{-C_1\pi})}{4b_K(e^{C_1\pi} - e^{-C_1\pi})}$$
$$+ \frac{\sqrt{4(1-\delta)b_K X_K M (e^{C_1\pi} - e^{-C_1\pi})^2 + \frac{1}{4}\left((b_K - d)X_K(e^{C_1\pi} - e^{-C_1\pi})(-2+\delta) + \delta C_2(e^{C_1\pi} + e^{-C_1\pi})\right)^2}}{2b_K(e^{C_1\pi} - e^{-C_1\pi})}.$$

This is non-negative only if $C_2 \in \mathbb{R}$ and $e^{C_1\pi} > 1$. If $C_2 \in \mathbb{R}$, then $C_1$ must also be real and positive, hence $e^{C_1\pi} > 1$ is automatically true and we need only check $C_2 \in \mathbb{R}$. This is true if:

$$4b_K M + (b_K - d)^2 X_K \geq 0,$$

which is clearly true. Thus $\hat{N}$ is a biologically-viable solution.

As before (Sec. S1.2), to study the stability of a periodic solution $\left(\hat{p}, \hat{N}\right)$, we consider the behaviour of a small perturbation to the solution: $(p(t), N(t)) = \left(\hat{p} + \xi_1(t), \hat{N} + \xi_2(t)\right)$. These perturbations may be written as:

$$\begin{pmatrix} \xi_1(t) \\ \xi_2(t) \end{pmatrix} = \Phi(t) \begin{pmatrix} \xi_1(0) \\ \xi_2(0) \end{pmatrix},$$

where $\Phi(t)$ is the matrix satisfying:

$$\frac{d\Phi}{dt} = \left.\begin{pmatrix} \frac{\partial}{\partial p}\tilde{f}_1(p,N) & \frac{\partial}{\partial N}\tilde{f}_1(p,N) \\ \frac{\partial}{\partial p}\tilde{f}_2(p,N) & \frac{\partial}{\partial N}\tilde{f}_2(p,N) \end{pmatrix}\right|_{p=\hat{p},N=\hat{N}} \Phi(t), \quad t \neq 2k\pi, \quad k = 0,1,2,\ldots, \quad \Phi(0) = \mathbf{I}.$$

The effect of our impulsive condition is given by:

$$\begin{pmatrix} \xi_1(2\pi) \\ \xi_2(2\pi) \end{pmatrix} = \begin{pmatrix} 1 & 0 \\ 0 & 1-\delta \end{pmatrix} \begin{pmatrix} \xi_1(2\pi^-) \\ \xi_2(2\pi^-) \end{pmatrix},$$

hence Floquet multipliers are the eigenvalues of:

$$M = \begin{pmatrix} 1 & 0 \\ 0 & 1-\delta \end{pmatrix} \Phi(2\pi).$$

Solving this for the $\hat{p} = 0$ solution, we get the two Floquet multipliers:

$$\rho_1 = \exp\left[\int_0^{2\pi} \left(b_r\left(1 - \frac{\hat{N}}{X_r}\right) - b_K\left(1 - \frac{\hat{N}}{X_K}\right) - \frac{M}{\hat{N}}\right) dt\right]$$
$$\rho_2 = (1-\delta)\exp\left[\int_0^{2\pi} \left(b_K\left(1 - 2\frac{\hat{N}}{X_K}\right) - d\right) dt\right].$$

Numerically, we see that $\rho_2$ is always less than 1, hence we need only check when $\rho_1 < 1$ for stability (see Mathematica file for details). The first Floquet multiplier is less than one only if the associated Floquet exponent is negative, where the exponent is given by Equation 5.

### *S2.2: Estimating the polymorphic solution*

When $\hat{p} = 0$ is not stable, coexistence of the two alleles arises so that $0 < \hat{p} < 1$. Since $p(t) = p(t^-)$, a long term periodic solution requires that the allele frequency at the beginning of the season is equal to the allele frequency at the end of the season. Therefore we assume the change in allele frequency throughout the season is small, hence:

$$\frac{dp}{dt} \approx 0.$$

Setting Eq. 4a equal to zero, we find:

$$p(N(t)) = \frac{b_r X_K N(N - X_r) - b_K X_r N(N - X_K) + X_r X_K M}{b_r X_K N(N - X_r) - b_K X_r N(N - X_K)}.$$

Plugging this in for $p$, Eq. 4b reduces to:

$$\frac{dN}{dt} = b_r N - \left( d + \frac{b_r}{X_r} N \right) N,$$

which along with our impulse restriction gives us the approximate periodic solution:

$$\hat{N}(t) = \frac{(b_r - d) X_r}{b_r} \frac{e^{-2\pi(b_r - d)} - (1 - \delta)}{e^{-2\pi(b_r - d)} - (1 - \delta) - \delta e^{-(b_r - d)t}}.$$

Plugging this into Eq. 4a, we get the following differential equation:

$$\frac{dp}{dt} = a_1(t) p + a_2(t) p^2,$$

where:

$$\begin{aligned} a_1(t) = (b_r - b_K) - \frac{b_r M}{(b_r - d) X_r} + \frac{b_r M \delta e^{-(b_r - d)t}}{(b_r - d) X_r \left( e^{-2\pi(b_r - d)} - (1 - \delta) \right)} \\ - \frac{(b_r - d)(b_r X_K - b_K X_r) \left( e^{-2\pi(b_r - d)} - (1 - \delta \right)}{b_r X_r \left( e^{-2\pi(b_r - d)} - (1 - \delta) - \delta e^{-(b_r - d)t} \right)}, \\ a_2(t) = -(b_r - b_K) + \frac{(b_r - d)(b_r X_K - b_K X_r) \left( e^{-2\pi(b_r - d)} - (1 - \delta \right)}{b_r X_r \left( e^{-2\pi(b_r - d)} - (1 - \delta) - \delta e^{-(b_r - d)t} \right)}. \end{aligned}$$

Using the change of variables (Vahidi et al., 2013):

$$p(t) = -\frac{y'(t)}{a_2(t) y(t)}, \quad y'(0) = -a_2(0) p(0), \quad y(0) = 1,$$

we derive an approximate long-term periodic solution $\hat{p}(t)$ for the polymorphic state:

$$\hat{p}(t) = \frac{C b(t)}{1 - C \int_0^t a_2(s) b(s) ds},$$

where:

$$\begin{aligned} b(t) = \exp \left[ \int a_1(t) \right] \\ C = -\frac{\hat{p}(0)}{b(0)}. \end{aligned}$$

To satisfy the requirements of a periodic solution, we require that $\hat{p}(0) = \hat{p}(2\pi)$ so that:

$$\hat{p}(0) = \frac{b(0) - b(2\pi)}{\int_0^{2\pi} a_2(s) b(s) ds}.$$

This solution cannot be written in closed form and must be computed numerically.
