## Supplementary material for "Local Adaptation of Life-History Traits in a Seasonal Environment": Mathematica Notebook PDF

In[343]:=

```
SetDirectory[NotebookDirectory[]];
```

### Floquet analysis for the island-mainland model

#### Model

In[344]:=

$$f1[p\_, n\_] := p\,(1-p)\left(bA\left(1-\frac{n}{XA}\right) - ba\left(1-\frac{n}{Xa}\right)\right) - p\,\frac{M}{n}$$

$$f2[p\_, n\_] := \left(bA\left(1-\frac{n}{XA}\right) p + ba\left(1-\frac{n}{Xa}\right)(1-p) - d\right) n + M$$

#### When $\hat{p} = 0$, solve for $\hat{N}(t)$

In[365]:=

```
temp = DSolve[{n'[t] == f2[0, n[t]], n[0] == n0}, n[t], t][[1]] // FullSimplify
```

Solve : Inverse functions are being used by Solve, so some solutions may not be found; use Reduce for complete solution information.

Out[365]=

$$\left\{n[t] \to \frac{1}{2\,ba}\left((ba-d)\,Xa + \sqrt{Xa}\,\sqrt{4\,ba\,M + (ba-d)^2\,Xa}\;\mathrm{Tanh}\left[\frac{t\,\sqrt{4\,ba\,M + (ba-d)^2\,Xa}}{2\,\sqrt{Xa}} + \mathrm{ArcTanh}\left[\frac{2\,ba\,n0 - ba\,Xa + d\,Xa}{\sqrt{Xa}\,\sqrt{4\,ba\,M + (ba-d)^2\,Xa}}\right]\right]\right)\right\}$$

Printed by Wolfram Mathematica Student Edition

In[368]:=

```
TrigToExp[n[t] /. temp] // FullSimplify
```

Out[368]=

$$\left(e^{\frac{t\sqrt{4\,ba\,M+(ba-d)^2\,Xa}}{\sqrt{Xa}}}\,(2\,M+(ba-d)\,n0)\,Xa+(-2\,M+(-ba+d)\,n0)\,Xa+n0\,\sqrt{Xa}\,\sqrt{4\,ba\,M+(ba-d)^2\,Xa}+e^{\frac{t\sqrt{4\,ba\,M+(ba-d)^2\,Xa}}{\sqrt{Xa}}}\,n0\,\sqrt{Xa}\,\sqrt{4\,ba\,M+(ba-d)^2\,Xa}\right)\Bigg/\left(d\left(-1+e^{\frac{t\sqrt{4\,ba\,M+(ba-d)^2\,Xa}}{\sqrt{Xa}}}\right)Xa-ba\left(-1+e^{\frac{t\sqrt{4\,ba\,M+(ba-d)^2\,Xa}}{\sqrt{Xa}}}\right)(-2\,n0+Xa)+\left(1+e^{\frac{t\sqrt{4\,ba\,M+(ba-d)^2\,Xa}}{\sqrt{Xa}}}\right)\sqrt{Xa}\,\sqrt{4\,ba\,M+(ba-d)^2\,Xa}\right)$$

Rewrite $\hat{N}$ in a simplified form:

$$(*c1=\frac{\sqrt{4\ ba\ M+(ba-d)^2\ Xa}}{\sqrt{Xa}}*)$$

$$(*c2=\sqrt{Xa}\ \sqrt{4\ ba\ M+(ba-d)^2\ Xa}*)$$

In[482]:=

$$nhatsimp[t\_] := \left(e^{c1\,t}\,(2\,M+(ba-d)\,n0)\,Xa+(-2\,M+(-ba+d)\,n0)\,Xa+n0\,c2+e^{c1\,t}\,n0\,c2\right)\Big/\left(d\left(-1+e^{c1\,t}\right)Xa-ba\left(-1+e^{c1\,t}\right)(-2\,n0+Xa)+\left(1+e^{c1\,t}\right)c2\right)\ //\ FullSimplify$$

Find $N_0$ such that $\hat{N}$ is periodic:

In[483]:=

```
n0sol = TrigToExp[FullSimplify[Solve[(1 - δ) nhatsimp[2 π] == nhatsimp[0], n0][[1]]]];
```

In[484]:=

$$nsol[t\_] := \left(e^{\frac{t\sqrt{4\,ba\,M+(ba-d)^2\,Xa}}{\sqrt{Xa}}}\,(2\,M+(ba-d)\,n0)\,Xa+(-2\,M+(-ba+d)\,n0)\,Xa+n0\,\sqrt{Xa}\,\sqrt{4\,ba\,M+(ba-d)^2\,Xa}+e^{\frac{t\sqrt{4\,ba\,M+(ba-d)^2\,Xa}}{\sqrt{Xa}}}\,n0\,\sqrt{Xa}\,\sqrt{4\,ba\,M+(ba-d)^2\,Xa}\right)\Bigg/\left(d\left(-1+e^{\frac{t\sqrt{4\,ba\,M+(ba-d)^2\,Xa}}{\sqrt{Xa}}}\right)Xa-ba\left(-1+e^{\frac{t\sqrt{4\,ba\,M+(ba-d)^2\,Xa}}{\sqrt{Xa}}}\right)(-2\,n0+Xa)+\left(1+e^{\frac{t\sqrt{4\,ba\,M+(ba-d)^2\,Xa}}{\sqrt{Xa}}}\right)\sqrt{Xa}\,\sqrt{4\,ba\,M+(ba-d)^2\,Xa}\right)\ /.\ n0sol\ /.$$

$$\left\{c1\to\frac{\sqrt{4\,ba\,M+(ba-d)^2\,Xa}}{\sqrt{Xa}},\ c2\to\sqrt{Xa}\,\sqrt{4\,ba\,M+(ba-d)^2\,Xa}\right\}$$

### Floquet Multiplier 1

The first Floquet multiplier is given by: $\rho_1 = \exp\left[\int_0^{2\pi}\left(b_A\left(1-\frac{\hat{N}}{X_A}\right)-b_a\left(1-\frac{\hat{N}}{X_a}\right)-\frac{M}{\hat{N}}\right)dt\right]$.

For this to be less than one, we need only that the exponent be negative, so we look only at the integral:

$\lambda_1 = \int_0^{2\pi}\left(b_A\left(1-\frac{\hat{N}}{X_A}\right)-b_a\left(1-\frac{\hat{N}}{X_a}\right)-\frac{M}{\hat{N}}\right)dt$

In[377]:=

```
lam1[pars_] :=
 NIntegrate[bA (1 - nsol[t]/XA) - ba (1 - nsol[t]/Xa) - M/nsol[t] /. pars, {t, 0, 2 π}]
```

In[378]:=

```
pars = {bA → 0.25, ba → 0.2, XA → 5000, Xa → 10 000, M → 1, δ → 0.65, d → 0.05};
```

In[379]:=

```
lam1[pars]
```

Out[379]=

```
0.188
```

#### Stability plot

Parameters: $b_A = 0.25, b_a = 0.2, X_A = 500, X_a = 1000, M = 1$

In[ ]:=

```
BA = 0.25;
Ba = 0.2;
coords = MeshCoordinates[DiscretizeRegion[
    ImplicitRegion[0 < d < Ba && 0 < delt < 1 - Exp[-2 π (BA - d)], {d, delt}],
    AccuracyGoal → 5, MaxCellMeasure → {"Length" → .01}]];

Export["coarse_coords.csv", coords, "CSV"];
```

In[ ]:=

```
temp = {#2, #, lam1[{bA → BA, ba → Ba, XA → 5000, Xa → 10 000, M → 1, d → #, δ → #2}]} & @@@
   coords;

Export["stability_lam1_data.csv", temp, "CSV"];
```

### Floquet Multiplier 2

The second Floquet multiplier is given by: $\rho_2 = (1-\delta)\exp\left[\int_0^{2\pi}\left(b_a - d - 2\frac{b_a}{X_a}\hat{N}\right)dt\right]$

In[387]:=

```
rho2[delt_, pars_] :=
 (1 - delt) Exp[Integrate[ba - d - 2 ba/Xa nsol[t] /. {δ → delt} /. pars, {t, 0, 2 π}]]
```

In[388]:=

```
Manipulate[rho2[δ, {ba → 0.4, Xa → 10 000, d → 0.2, M → 1}], {δ, 0, 1}]
```

Out[388]=

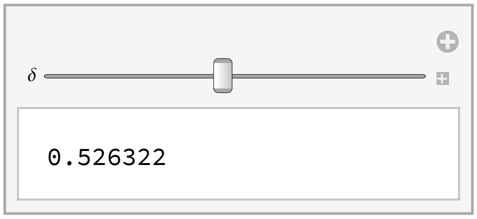


In[389]:=

```
Manipulate[rho2[δ, {ba → 0.2, Xa → 10 000, d → 0.05, M → 0.5}], {δ, 0, 1}]
```

Out[389]=

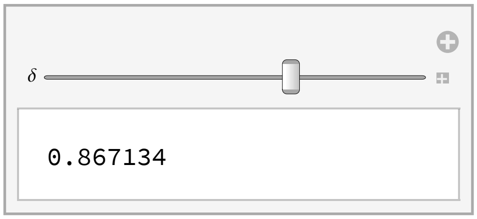


In[∘]:=

```
temp2 = {#2, #, rho2[#2, {ba → 0.2, Xa → 10 000, d → #, M → 1}]} & @@@ coords;
```

In[∘]:=

```
Export["stability_rho2_data.csv", temp2, "CSV"];
```

In[392]:=

```
ListContourPlot[temp2, PlotLegends → Automatic,
 RegionFunction → Function[{x, y, z}, 1 - Exp[-2 π (BA - y)] > x && z < 1],
 Frame → {{True, True}, {True, True}},
 FrameStyle → Directive[Black],
 FrameLabel → {Style["δ", 15], Style["d", 15]},
 ColorFunction → ColorData["GrayTones"]]
```

Out[392]=

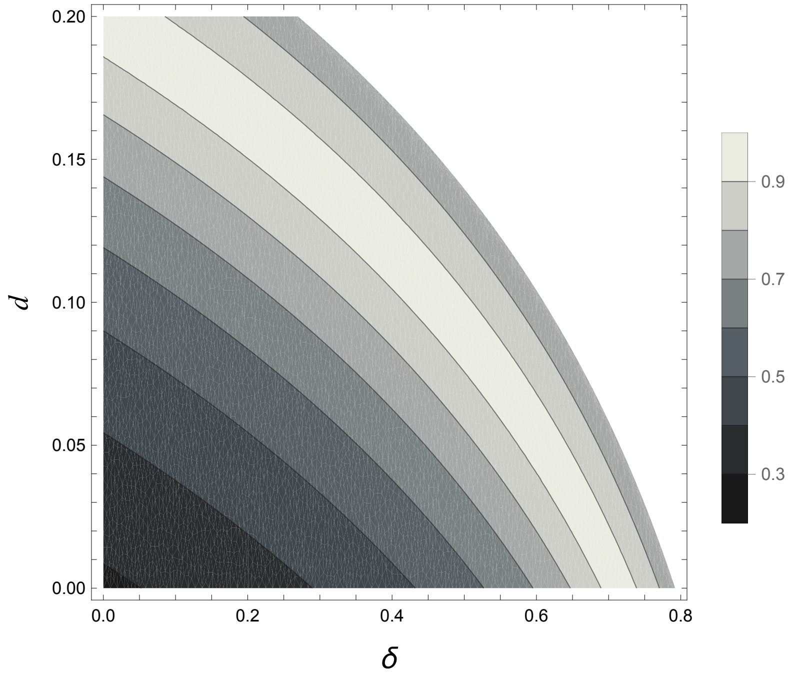


Numerically, we see that $\rho_2 < 1$.

### Get region of stability

In[400]:=

```
data = Import["stability_lam1_data.csv"];
```

In[401]:=

```
temp = ListContourPlot[data, InterpolationOrder → 2,
   Contours → {0},
   ContourShading → None,
   ContourStyle → Directive[Thick, Red],
   Frame → {{True, True}, {True, True}},
   FrameTicks → {{True, False}, {True, False}},
   FrameLabel → {Style["d", 15], Style["δ", 15]},
   FrameStyle → Directive[Black]];
```

In[402]:=
```
line1 = First@Cases[Normal[temp], _Line, Infinity];
line2 = Last@Cases[Normal[temp], _Line, Infinity];
```

In[404]:=
```
tval1 = ResourceFunction["LeeInterpolatingNodes"][line1[[1]]];
tval2 = ResourceFunction["LeeInterpolatingNodes"][line2[[1]]];
```

In[406]:=
```
cc1 = Interpolation[Transpose[{tval1, line1[[1]]}], Method → "Spline"];
cc2 = Interpolation[Transpose[{tval2, line2[[1]]}], Method → "Spline"];
```

In[409]:=
```
lin1 = Table[cc1[s], {s, 0, 1.03, 0.01}];
lin2 = Table[cc2[s], {s, 0, 1.03, 0.01}];
```

In[412]:=
```
mr = MeshRegion[DiscretizeRegion[Region[Polygon[Join[lin2, lin1]]]]];
```

### Polymorphism approximation for island-mainland model

#### Assume $\frac{dp}{dt} \approx 0$

Solve for $p(N)$

In[ ]:=
```
Solve[f1[p, n] == 0, p]
```

Out[ ]=

$$\left\{\{p \to 0\},\ \left\{p \to \frac{bA\,n^2\,Xa - ba\,n^2\,XA + M\,Xa\,XA + ba\,n\,Xa\,XA - bA\,n\,Xa\,XA}{n\,(bA\,n\,Xa - ba\,n\,XA + ba\,Xa\,XA - bA\,Xa\,XA)}\right\}\right\}$$

#### Approximate $\hat{N}(t)$

Plug $p(N)$ into $f_2(p, N)$ and solve the resulting differential equation for $\frac{dN}{dt} = f_2\,(p\,(N), N)$ to get $\hat{N}(t)$.

In[•]:=
```
f2[p, n] /. {p → (bA n^2 Xa - ba n^2 XA + M Xa XA + ba n Xa XA - bA n Xa XA)/(n (bA n Xa - ba n XA + ba Xa XA - bA Xa XA))} // Simplify
```
Out[•]=

$$n\left(bA - d - \frac{bA\,n}{XA}\right)$$

In[•]:=
```
DSolve[{n'[t] == n[t] (bA - d - bA n[t]/XA), n[0] == n0}, n[t], t] // FullSimplify
```

••• Solve : Inverse functions are being used by Solve, so some solutions may not be found; use Reduce for complete solution information.

Out[•]=

$$\left\{\left\{n[t] \to \frac{(bA - d)\,n0\,XA}{bA\,n0 - e^{(-bA+d)\,t}\,(bA\,(n0 - XA) + d\,XA)}\right\}\right\}$$

Solve for $\hat{N}(0) = N_0$ based on the requirement that $(1-\delta)\hat{N}(2\pi) = N_0$

In[•]:=
```
Solve[((1 - δ) (bA - d) n0 XA)/(bA n0 - e^((-bA+d) 2 π) (bA (n0 - XA) + d XA)) == n0, n0] // Simplify
```
Out[•]=

$$\left\{\{n0 \to 0\},\ \left\{n0 \to \frac{(bA - d)\,XA\,\left(-1 + e^{2\,(-bA+d)\,\pi} + \delta\right)}{bA\,\left(-1 + e^{2\,(-bA+d)\,\pi}\right)}\right\}\right\}$$

In[451]:=
```
N0 = ((bA - d) XA (-1 + e^(2 (-bA+d) π) + δ))/(bA (-1 + e^(2 (-bA+d) π))) // Simplify;
```

In[452]:=
```
Nf = N0/(1 - δ) // Simplify;
```

Thus $\hat{N}(t)$ is given by:

In[395]:=
```
((bA - d) n0 XA)/(bA n0 - e^((-bA+d) t) (bA (n0 - XA) + d XA)) /.
  {n0 → ((bA - d) XA (-1 + e^(2 (-bA+d) π) + δ))/(bA (-1 + e^(2 (-bA+d) π)))} // FullSimplify
```
Out[395]=

$$\frac{(bA - d)\,XA\,\left(-1 + e^{2\,(-bA+d)\,\pi} + \delta\right)}{bA\,\left(-1 + e^{2\,(-bA+d)\,\pi} + \delta - e^{(-bA+d)\,t}\,\delta\right)}$$

Determine when $\hat{N}(t)$ is a biologically valid solution, that it, when $\hat{N}(t) \geq 0$.

In[∘]:=
```
Reduce[{((bA - d) XA (-1 + e^(2 (-bA+d) π) + δ))/(bA (-1 + e^(2 (-bA+d) π))) > 0, 0 < δ < 1, XA > 0, bA > d > 0}]
```

Out[∘]=

$bA > 0 \&\& 0 < d < bA \&\& XA > 0 \&\& 0 < \delta < 1 - e^{2(-bA+d)\pi}$

This is the same as the criteria for a population of exclusively $A$ allele individuals to be viable.

### Approximate $\hat{p}(t)$

Plug $\hat{N}(t)$ back into $f_1(p, N)$ and solve the resulting differential equation $\frac{dp}{dt} = f_1(p, \hat{N})$ to get $\hat{p}(t)$.

$\frac{dp}{dt} = f_1(p, \hat{N})$ takes the form $\frac{dp}{dt} = a_1(t)\,p + a_2(t)\,p^2$, where $a_1(t)$ and $a_2(t)$ are given by:

In[∘]:=
```
CoefficientList[f1[p, n] /. {n → ((bA - d) XA (-1 + e^(2 (-bA+d) π) + δ))/(bA (-1 + e^(2 (-bA+d) π) + δ - e^((-bA+d) t) δ))}, p] // FullSimplify
```

Out[∘]=

$$\left\{0,\ -ba + bA - \frac{bA\,M}{bA\,XA - d\,XA} + \frac{bA\,e^{(-bA+d)\,t}\,M\,\delta}{(bA-d)\,XA\left(-1 + e^{2(-bA+d)\pi} + \delta\right)} - \frac{(bA-d)\,(bA\,Xa - ba\,XA)\left(-1 + e^{2(-bA+d)\pi} + \delta\right)}{bA\,Xa\left(-1 + e^{2(-bA+d)\pi} + \delta - e^{(-bA+d)\,t}\,\delta\right)},\ ba - bA + \frac{(bA-d)\,(bA\,Xa - ba\,XA)\left(-1 + e^{2(-bA+d)\pi} + \delta\right)}{bA\,Xa\left(-1 + e^{2(-bA+d)\pi} + \delta - e^{(-bA+d)\,t}\,\delta\right)}\right\}$$

In[396]:=

$$a1[t\_] := -ba + bA - \frac{bA\,M}{bA\,XA - d\,XA} + \frac{bA\,e^{(-bA+d)\,t}\,M\,\delta}{(bA-d)\,XA\left(-1 + e^{2(-bA+d)\pi} + \delta\right)} - \frac{(bA-d)\,(bA\,Xa - ba\,XA)\left(-1 + e^{2(-bA+d)\pi} + \delta\right)}{bA\,Xa\left(-1 + e^{2(-bA+d)\pi} + \delta - e^{(-bA+d)\,t}\,\delta\right)}$$

$$a2[t\_] := ba - bA + \frac{(bA-d)\,(bA\,Xa - ba\,XA)\left(-1 + e^{2(-bA+d)\pi} + \delta\right)}{bA\,Xa\left(-1 + e^{2(-bA+d)\pi} + \delta - e^{(-bA+d)\,t}\,\delta\right)}$$

We know that $\hat{p}(t) = \frac{C\,b(t)}{1 - C\int_0^t a_2(s)\,b(s)\,ds}$, where $b(t) = \exp\left[\int a_1(t)\right]$ and $C = -\frac{\hat{p}(0)}{b(0)}$

In[ ]:=
```
Integrate[a1[t], t] // FullSimplify
```
Out[ ]=

$$-\text{ba}\,t+\text{bA}\left(t-\frac{M\,t}{\text{bA XA}-d\,\text{XA}}-\frac{e^{2\,\text{bA}\,\pi-\text{bA}\,t+d\,t}\,M\,\delta}{(\text{bA}-d)^2\,\text{XA}\left(e^{2\,d\,\pi}+e^{2\,\text{bA}\,\pi}\,(-1+\delta)\right)}\right)+\left(-1+\frac{\text{ba XA}}{\text{bA Xa}}\right)\text{Log}\left[e^{(\text{bA}-d)\,t}+e^{(\text{bA}-d)\,(2\,\pi+t)}\,(-1+\delta)-e^{2\,(\text{bA}-d)\,\pi}\,\delta\right]$$

In[ ]:=
```
Exp[-ba t + bA (t - M t/(bA XA - d XA) - (e^(2 bA π - bA t + d t) M δ)/((bA - d)^2 XA (e^(2 d π) + e^(2 bA π) (-1 + δ)))) +
  (-1 + (ba XA)/(bA Xa)) Log[e^((bA-d) t) + e^((bA-d) (2 π+t)) (-1 + δ) - e^(2 (bA-d) π) δ]] // FullSimplify
```
Out[ ]=

$$e^{-\text{ba}\,t+\text{bA}\left(t-\frac{M\,t}{\text{bA XA}-d\,\text{XA}}-\frac{e^{2\,\text{bA}\,\pi-\text{bA}\,t+d\,t}\,M\,\delta}{(\text{bA}-d)^2\,\text{XA}\left(e^{2\,d\,\pi}+e^{2\,\text{bA}\,\pi}\,(-1+\delta)\right)}\right)}\left(e^{(\text{bA}-d)\,t}+e^{(\text{bA}-d)\,(2\,\pi+t)}\,(-1+\delta)-e^{2\,(\text{bA}-d)\,\pi}\,\delta\right)^{-1+\frac{\text{ba XA}}{\text{bA Xa}}}$$

In[424]:=
```
b[t_] := e^(-ba t+bA (t - M t/(bA XA-d XA) - (e^(2 bA π-bA t+d t) M δ)/((bA-d)^2 XA (e^(2 d π)+e^(2 bA π) (-1+δ))))) (e^((bA-d) t) + e^((bA-d) (2 π+t)) (-1 + δ) - e^(2 (bA-d) π) δ)^(-1+(ba XA)/(bA Xa))
```

Setting $\hat{p}(0)=\hat{p}(2\pi)$, we solve for $\hat{p}(0)$:

In[ ]:=
```
Solve[(-(p0/b0) b2π)/(1 + (p0/b0) int) == p0, p0]
```
Out[ ]=

$$\left\{\{p0\to 0\},\left\{p0\to\frac{-b0-b2\pi}{\text{int}}\right\}\right\}$$

Hence $\hat{p}(0)=\hat{p}(2\pi)=\dfrac{b(0)-b(2\pi)}{\int_0^{2\pi}a_2(s)\,b(s)\,ds}$.

In[421]:=
```
pars = {bA → 0.25, ba → 0.2, XA → 500, Xa → 1000, M → 1, δ → 0.65, d → 0.05};
```
In[422]:=
```
p0[pars_] := Re[(b[0] - b[2 π] /. pars) / NIntegrate[a2[s] × b[s] /. pars, {s, 0, 2 π}]]
```
In[423]:=
```
p0[pars]
```
Out[423]=
```
0.451231
```

#### Calculate frequencies

Parameters: $b_A=0.25, b_a=0.2, X_A=500, X_a=1000, M=1$

In[459]:=

```
BA = 0.25;
Ba = 0.2;
coords = MeshCoordinates[DiscretizeRegion[
    ImplicitRegion[0 < d < Ba && 0 < delt < 1 - Exp[-2 π (BA - d)], {d, delt}],
    AccuracyGoal → 5, MaxCellMeasure → {"Length" → .005}]];
```

In[420]:=

```
Export["fine_coords.csv", coords, "CSV"];
```

In[462]:=

```
temp2 = {#2, #, If[{#2, #} ∈ mr, p0[
       {bA → BA, ba → Ba, XA → 5000, Xa → 10 000, M → 1, δ → #2, d → #}], 0]} & @@@ coords;
```

In[463]:=

```
Export["frequency_data.csv", temp2, "CSV"];
```

### Measuring local adaptation

We define local adaptation generally by:

$\Delta_{LF} = E[W_{ii}] - E_j[W_{ij}]$

Which is the difference between the expected fitness of the population of interest in its own habitat and its expected fitness in all possible fitnesses.

In the case of the island-mainland model, this becomes:

$$\Delta_{LF} = \overline{W}_{I\to I} - \frac{1}{2}\left(\overline{W}_{I\to I} + \overline{W}_{I\to M}\right) = \frac{1}{2}\,\overline{W}_{I\to I} - \frac{1}{2}\,\overline{W}_{I\to M}$$

$$= \frac{1}{2}\left(p\,W_A + (1-p)\,W_a\right) - \frac{1}{2}\,W_a$$

Fitness is given by:

$$W_i = b_i - \left(d + \frac{N}{X_i}\right).$$

Note that since $p(t)$ and $N(t)$ are time dependent, there are different ways to define local adaptation throughout the cycle. We start by considering the local adaptation at the beginning and end of the cycle.

In[414]:=

```
WA[N_, pars_] := bA - (d + N bA / XA) /. pars
Wa[N_, pars_] := ba - (d + N ba / Xa) /. pars
```

In[416]:=

```
LF[p_, N_, pars_] := (p / 2) (WA[N, pars] - Wa[N, pars])
```

In[466]:=

```
frequencies = Import["frequency_data.csv"];
```

In[467]:=

```
LFinit = {#, #2, If[{#, #2} ∈ mr,
      LF[#3, N0 /. {bA → BA, ba → Ba, XA → 5000, Xa → 10 000, M → 1, δ → #, d → #2},
       {bA → BA, ba → Ba, XA → 5000, Xa → 10 000, d → #2}], 0]} & @@@ frequencies;
```

In[468]:=

```
Export["local_adaptation_init.csv", LFinit, "CSV"];
```

In[469]:=

```
LFfinal = {#, #2, If[{#, #2} ∈ mr,
      LF[#3, Nf /. {bA → BA, ba → Ba, XA → 5000, Xa → 10 000, M → 1, δ → #, d → #2},
       {bA → BA, ba → Ba, XA → 5000, Xa → 10 000, d → #2}], 0]} & @@@ frequencies;
```

In[470]:=

```
Export["local_adaptation_final.csv", LFfinal, "CSV"];
```

#### Stochastic data

Parameters: $b_A = 0.25, b_a = 0.2, X_A = 500, X_a = 1000, M = 1$

```
BA = 0.25;
Ba = 0.2;
xA = 5000;
xa = 10 000;
```

In[517]:=

```
temp = Import["stoch_data.csv"];
```

In[523]:=

```
stodata = Drop[temp, 1, 2];
```

In[527]:=

```
stolf0 = {#2, #, Chop[LF[Chop[#3], #4, {bA → BA, ba → Ba, XA → xA, Xa → xa, d → #}]]} & @@@
   stodata;
```

In[529]:=

```
Export["stoch_local_adaptation_init.csv", stolf0, "CSV"];
```

In[531]:=

```
stolff = {#2, #, Chop[LF[Chop[#5], #6, {bA → BA, ba → Ba, XA → xA, Xa → xa, d → #}]]} & @@@
   stodata;
```

In[532]:=

```
Export["stoch_local_adaptation_final.csv", stolff, "CSV"];
```

### Measuring strength of genetic drift

In[476]:=

```
Scoeff[n_, pars_] := WA[n, pars] - Wa[n, pars]
```

#### When p > 0:

In[503]:=

```
nhat[t_] := (bA - d) n0 XA / (bA n0 - e^((-bA+d) t) (bA (n0 - XA) + d XA)) /. {n0 → N0}
```

In[506]:=

```
p1neff[pars_] := 2 π / NIntegrate[1/(nhat[t]/.pars), {t, 0, 2 π}]
```

#### When p = 0:

In[489]:=

```
p0neff[pars_] := 2 π / NIntegrate[1/(nsol[t]/.pars), {t, 0, 2 π}]
```

Parameters: $b_A = 0.25, b_a = 0.2, X_A = 500, X_a = 1000, M = 1$

In[490]:=

```
BA = 0.25;
Ba = 0.2;
xA = 5000;
xa = 10 000;
m = 1;
```

In[507]:=

```
neffdata = {#2, #,
      If[{#2, #} ∈ mr, p1neff[{bA → BA, ba → Ba, XA → xA, Xa → xa, M → m, δ → #2, d → #}],
       p0neff[{bA → BA, ba → Ba, XA → xA, Xa → xa, M → m, δ → #2, d → #}]]} &@@@ coords;
```

In[508]:=

```
nesdata = {#, #2, Abs[Scoeff[#3, {bA → BA, ba → Ba, XA → xA, Xa → xa}]] #3} &@@@ neffdata;
```

In[511]:=

```
Export["nes_data.csv", nesdata, "CSV"];
```
